# Linking the *Salmonella enterica* 1,2-propanediol utilization bacterial microcompartment shell to the enzymatic core via the shell protein PduB

**DOI:** 10.1101/2021.10.27.466122

**Authors:** Nolan W. Kennedy, Carolyn E. Mills, Charlotte H. Abrahamson, Andre Archer, Michael C. Jewett, Niall M. Mangan, Danielle Tullman-Ercek

## Abstract

Bacterial microcompartments (MCPs) are protein-based organelles that house the enzymatic machinery for metabolism of niche carbon sources, allowing enteric pathogens to outcompete native microbiota during host colonization. While much progress has been made toward understanding MCP biogenesis, questions still remain regarding the mechanism by which core MCP enzymes are enveloped within the MCP protein shell. Here we explore the hypothesis that the shell protein PduB is responsible for linking the shell of the 1,2-propanediol utilization (Pdu) MCP from *Salmonella enterica* serovar Typhimurium LT2 to its enzymatic core. Using fluorescent reporters, we demonstrate that all members of the Pdu enzymatic core are encapsulated in Pdu MCPs. We also demonstrate that PduB is the sole protein responsible for linking the entire Pdu enzyme core to the MCP shell. Using MCP purifications, transmission electron microscopy, and fluorescence microscopy we find that shell assembly can be decoupled from the enzymatic core, as apparently empty MCPs are formed in *Salmonella* strains lacking PduB. Mutagenesis studies also reveal that PduB is incorporated into the Pdu MCP shell via a conserved, lysine-mediated hydrogen bonding mechanism. Finally, growth assays and systems-level pathway modeling reveal that unencapsulated pathway performance is strongly impacted by enzyme concentration, highlighting the importance of minimizing polar effects when conducting these functional assays. Together, these results provide insight into the mechanism of enzyme encapsulation within Pdu MCPs and demonstrate that the process of enzyme encapsulation and shell assembly are separate processes in this system, a finding that will aid future efforts to understand MCP biogenesis.

**Importance:** MCPs are unique, genetically encoded organelles used by many bacteria to survive in resource-limited environments. There is significant interest in understanding the biogenesis and function of these organelles, both as potential antibiotic targets in enteric pathogens and also as useful tools for overcoming metabolic engineering bottlenecks. However, the mechanism by which these organelles are formed natively is still not completely understood. Here we provide evidence of a potential mechanism in*S. enterica* by which a single protein, PduB, links the MCP shell and metabolic core. This finding is critical for those seeking to disrupt MCPs during pathogenic infections or for those seeking to harness MCPs as nanobioreactors in industrial settings.

## Introduction

Subcellular compartmentalization into organelles is an important feature of cellular life (1). Organelles allow cells to organize chemical transformations in space, enabling diverse metabolic processes to occur simultaneously within single cells. This functionality, sequestering diverse processes into membrane delimited organelles, is most frequently associated with the eukaryotic domain (1). However, prokaryotes have diverse modes of subcellular compartmentalization as well, including both lipid (2) and protein (3) bound organelles.

A prime example of a protein-delimited organelle is the bacterial microcompartment (MCP) (4). These protein-based organelles come in numerous varieties, including carbon-fixing carboxysomes found in cyanobacteria and carbon-harvesting metabolosomes often found in enteric pathogens (Figure 1A) (5, 6). Metabolosomes consist of a proteinaceous shell surrounding an encapsulated enzymatic core (Figure 1B) (6). Typically the enzymatic core consists of a specific metabolic pathway that utilizes a niche carbon source (Figure 1B), often one that must move through a toxic aldehyde intermediate (Figure 1B) (7). The MCP shell is thought to provide a diffusional barrier between the cytosol and enzymatic core, which both helps to increase flux through the encapsulated pathway and also protects the cell from the toxic intermediate (8, 9). In this way, MCPs are thought to be important for providing a growth advantage to pathogenic microbes during propagation within hosts (10).

**Figure 1.**
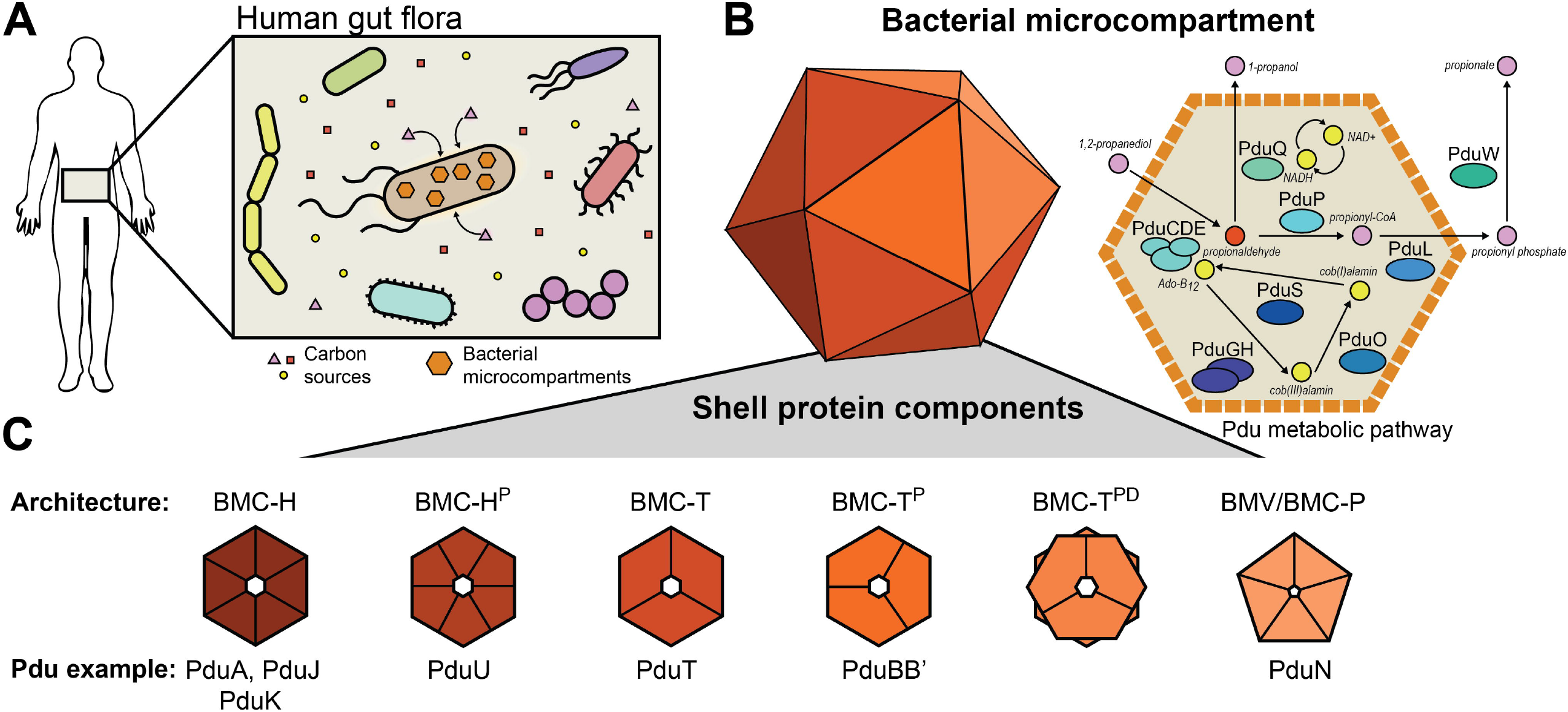
Schematic representation of cells containing Pdu MCPs. (A) Cells harboring MCPs can scavenge niche carbon sources from their environment. (B) The Pdu MCP encapsulates enzymes responsible for the metabolism of 1,2-propanediol to propionate and 1-propanol via the toxic intermediate propionaldehyde, which is sequestered by the MCP shell. The Pdu MCP also encapsulates various enzymes responsible for cofactor recycling. (C) MCPs are composed of a protein shell made of a variety of different shell protein architectures, including hexamers (BMC-H), trimers (BMC-T), and pentamers (BMC-P) and their various subtypes.

One of the most well-studied types of metabolosomes is the 1,2-propanediol utilization (Pdu) MCP found in *Salmonella enterica* serovar Typhimurium LT2 (hereafter referred to as “LT2”). The Pdu MCP provides an excellent model for metabolosome study as Pdu MCPs natively exist in a number of model bacterial hosts such as the aforementioned LT2 as well as strains of *Escherichia coli* (11). Pdu MCPs are broadly distributed throughout many bacterial phyla and numerous Pdu subtypes exist (12–14). The LT2 Pdu MCP is also a valuable model metabolosome because it contains representative shell proteins of all but one of the known shell protein subtypes (Figure 1C), making Pdu-based studies broadly applicable to a variety of MCP systems (13).

Generally, Pdu MCP shells are irregular and polyhedral in geometry and roughly 140 nm in diameter (15). They are composed of small, assembled shell proteins which tile together around the enzymatic core into a perforated, mosaic surface, containing small pores to allow metabolite diffusion (Figure 1B-C) (16). The assembled shell proteins come in a number of basic archetypal categories (Figure 1C) (13). These varying architectures include six-sided hexameric or pseudohexameric proteins composed of pfam00936 domains, as well as five-sided pentameric proteins (BMC-P or BMV) composed of pfam03319 domains (Figure 1C) (17, 18). The hexagonal pfam00936 proteins can be further subdivided into simple hexamers (BMC-H), circularly permutated hexamers (BMC-H^P^), pseudohexameric trimers (BMC-T), circularly permutated pseudohexameric trimers (BMC-T^P^), and circularly permutated pseudohexameric trimer dimers (BMC-T^DP^) (Figure 1C) (13). The LT2 Pdu operon contains eight total shell proteins, with representatives from all but the BMC-T^DP^ subcategory (19).

Although tremendous progress has been made toward understanding the biogenesis and assembly of the Pdu MCP, questions remain regarding the mechanism linking the shell and the enzymatic core. Numerous studies exist throughout the literature proposing nearly every single Pdu shell protein as responsible for encapsulating one or more of the enzymes that localize to the MCP core (Figure 2A) (20–24). In this study we expand on the hypothesis set forth by Lehman et al., which posited that the BMC-T^P^ protein PduB is responsible enzyme encapsulation in Pdu MCPs (20). To build on this study, we created a suite of fluorescently-tagged enzyme reporters which report on the encapsulation state of all enzyme interaction partners (i.e. at least one enzyme from sets of enzymes that are known to interact) (Figure 2B). This reveals that PduB is responsible for encapsulating all members of the enzymatic core. Surprisingly, we demonstrate that PduW, an enzyme previously thought to be cytosolic (25), is also encapsulated within MCPs. We also show that signal sequences are sufficient for targeting to the enzymatic core, indicating that signal sequences interact with other members of the enzymatic core and not necessarily the MCP shell as previously thought (21, 23). Interestingly, MCP purifications from strains with the PduB open reading frame knocked out demonstrate that the MCP shell still assembles in the absence of the enzymatic core. This provides the first conclusive evidence that shell assembly and enzymatic core assembly are separate processes that can be decoupled. Through mutagenesis, we show that PduB is incorporated into the MCP shell via a conserved hydrogen bonding mechanism, similar to the mechanism for incorporation of the BMC-H proteins PduA and PduJ (26–28). Finally, our work shows decoupling the enzymatic core from the MCP shell results in a surprising growth phenotype, differing in growth rate and metabolic profile from other disrupted MCPs. Together, these results provide clarity to a field crowded with conflicting evidence regarding MCP enzyme encapsulation and organelle biogenesis.

**Figure 2.**
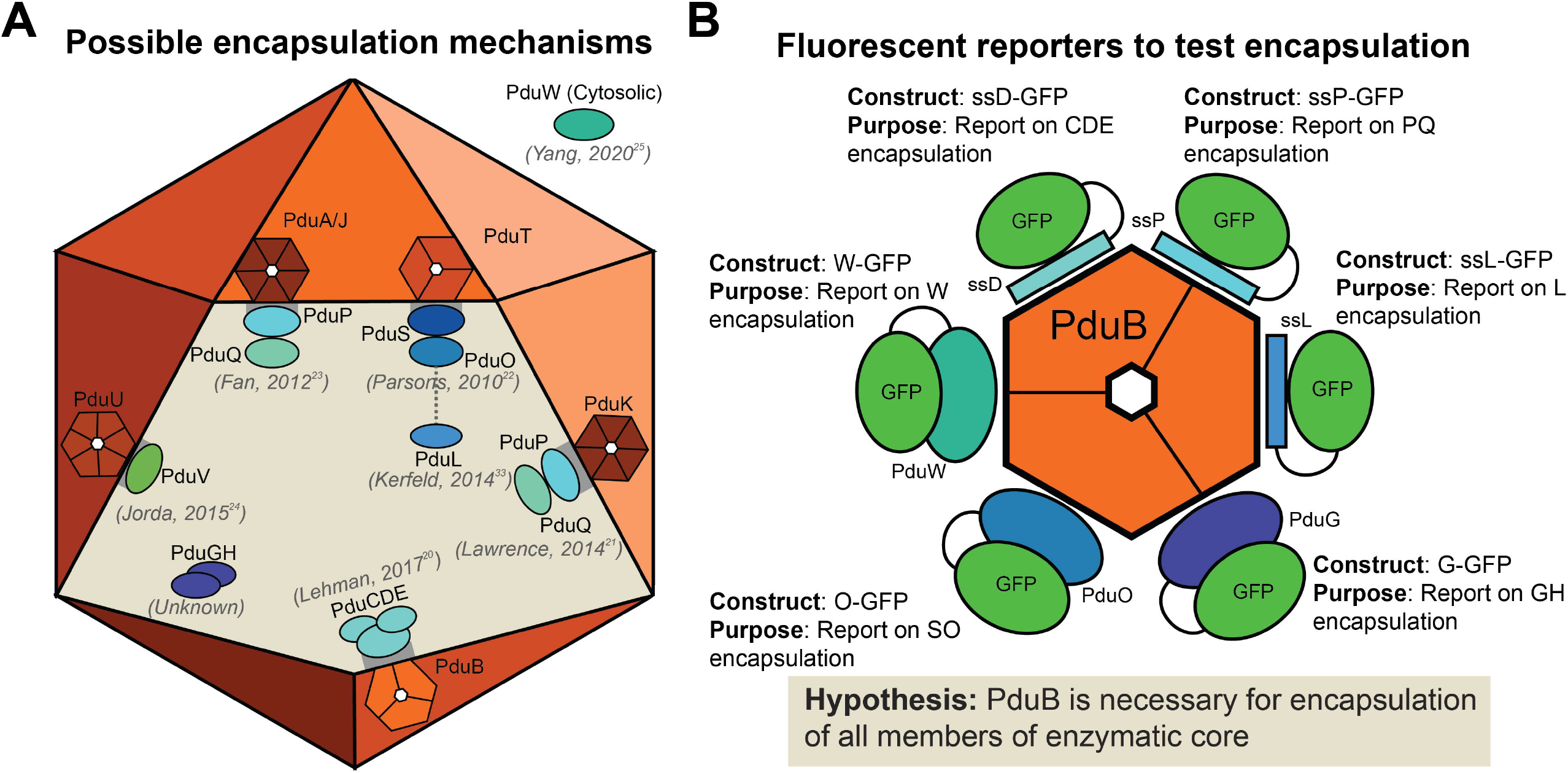
A suite of fluorescent reporters is used to investigate hypotheses about enzyme encapsulation within Pdu MCPs. (A) A number of hypotheses have been proposed in the Pdu MCP literature regarding the mechanism of enzyme encapsulation (20–25, 33). Six of the eight Pdu shell proteins have been suggested to have a role in enzyme encapsulation. In order to test the hypothesis proposed in Lehman et al., we created a suite of fluorescent reporters (B) to inform on the encapsulation state on the entire Pdu MCP core.

## Results

### Fluorescent reporters demonstrate PduB is critical for enzyme encapsulation

In 2017, Lehman et al. reported that deletion of the PduB N-terminus reduced the apparent enzyme content of purified MCPs as indicated by sodium dodecyl sulphate-polyacrylamide gel electrophoresis (SDS-PAGE). While groundbreaking, this study did not directly demonstrate effects of PduB deletion on enzyme localization to MCPs, as malformed MCPs or mistargeted enzymes are co-purified with MCPs in the differential centrifugation process (27). To build on this landmark study, we created a suite of green fluorescent protein (GFP) reporters to inform on which members of the enzymatic core are directly impacted by the loss of PduB. Numerous studies have indicated that many Pdu shell proteins directly bind and encapsulate various core enzymes. Thus, we reasoned that PduB may only be responsible for binding and encapsulating a select subset of Pdu enzymes. For example, studies have suggested that PduK (21), PduT (22), PduA (23), PduJ (23), and PduU (24) are responsible for binding and targeting one or more Pdu enzymes to the MCP core (Figure 2A). However, the work by Lehman et al. is the only study to date to directly show that a genetic alteration (deletion of the PduB N-terminus in the *pdu* operon) leads to changes in enzyme localization. We therefore hypothesized that PduB is the most likely shell protein to be responsible for enzyme targeting (20).

To test the hypothesis that PduB is responsible for linking the Pdu MCP shell and enzymatic core, we created GFP protein fusions to report on each Pdu enzyme (Figure 1B, 2B). Some enzymes or sets of enzymes contain known, short, N-terminal signal sequences that are sufficient for targeting enzymes to the MCP lumen. The signal sequence of PduD (ssD) is sufficient for targeting PduC, PduD, and PduE (29); the signal sequence for PduP (ssP) is sufficient for targeting PduP (30); and the signal sequence for PduL (ssL) is sufficient for targeting PduL (31). Since these are the minimal components necessary for targeting these proteins, they were fused to GFP and used to report on the enzyme’s change in encapsulation as a result of genetic manipulations in this study (Figure 2B). Since PduD interacts with PduC and PduE, only ssD-GFP was necessary to report on these three proteins. In the same way, since PduP interacts with PduQ (32), ssP-GFP was used to report on this enzyme set (PduP and PduQ). PduL has been hypothesized to interact with PduS (33), but this has not been confirmed experimentally. However, PduS interacts with PduO (34), and since PduO is the more abundant member of this pair (25), it was fused to GFP and used as a reporter for PduS and PduO (O-GFP) (Figure 2B). Similarly, PduG and PduH are subunits of a diol dehydratase reactivase enzyme and PduG is the more abundant member of this protein pair, so it was fused with GFP (G-GFP) and used for this study (Figure 2B) (25, 35–37). Finally, PduW, which is thought to be cytosolic (25), was fused C-terminally with GFP (W-GFP) to report on its encapsulation state (Figure 2B).

To test the hypothesis that PduB is necessary for the encapsulation of all members of the enzymatic core listed above, each fluorescent reporter was expressed in four strains of LT2: (1) wild type (WT) LT2, (2) a strain with the genes encoding the BMC-H proteins PduA and PduJ knocked out (ΔA ΔJ), (3) a strain with the transcriptional activator PocR knocked out (ΔpocR), and (4) an experimental strain with PduB knocked out (ΔB) to inform on the effect of PduB loss on enzyme encapsulation (Figure 3A). The WT strain serves as a positive control and enables visualization of be expressed from the genome in the presence of the native inducer 1,2-propanediol (1,2-PD) in this strain (Figure 3A) (27, 38–41). This allows us to determine which GFP- tagged constructs formed aggregates when expressed in the absence of MCPs and to what degree.

**Figure 3.**
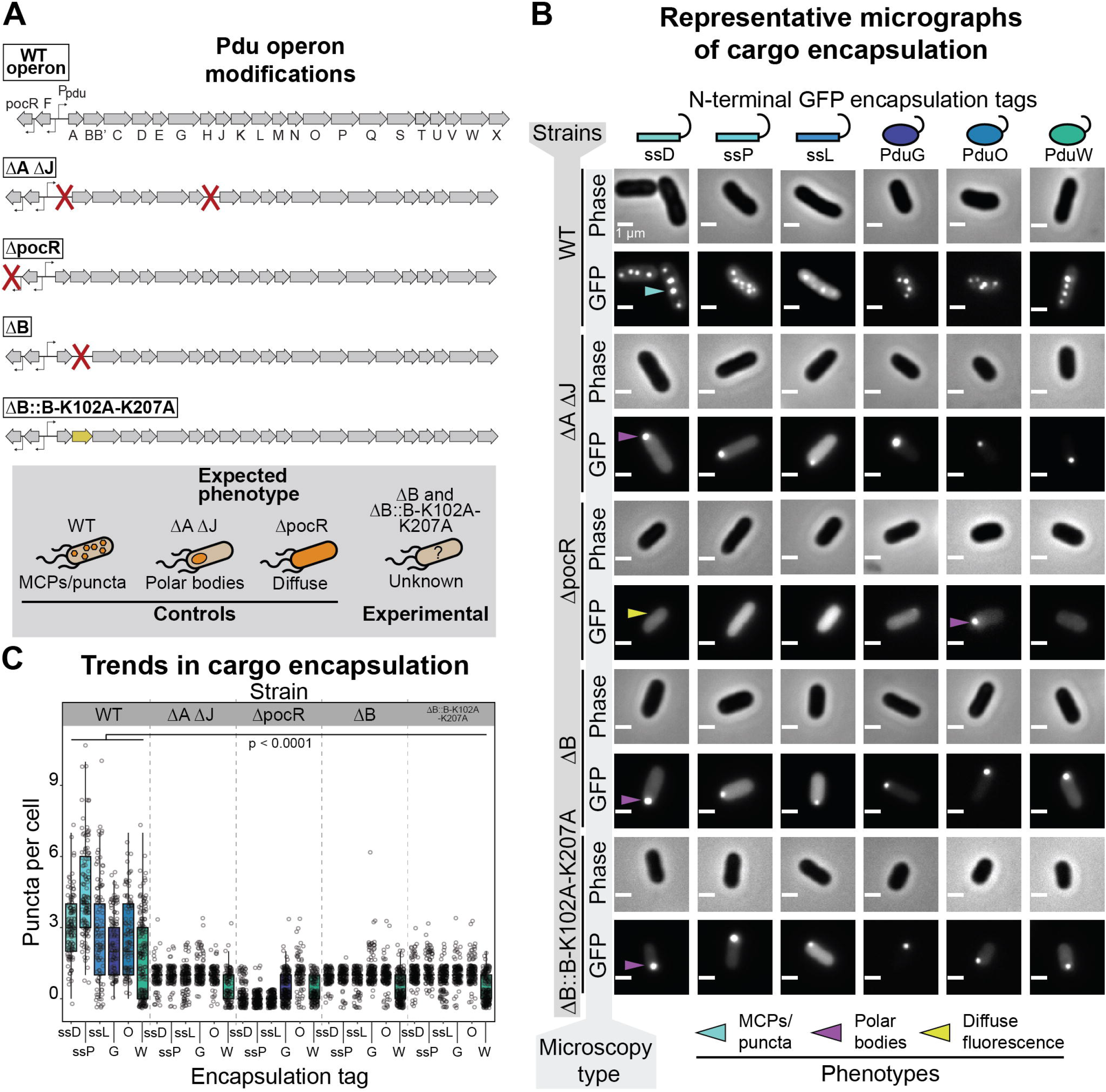
Loss of PduB disrupts enzyme encapsulation. (A) Schematic representation of the *pdu* operon and modifications to various LT2 strains used in this study, including the wild type (WT) operon. ΔA ΔJ has the open reading frames encoding PduA and PduJ knocked out, disrupting MCP formation. The ΔpocR strain has the transcriptional activator for the *pdu* operon knocked out, disrupting Pdu MCP expression. The ΔB strain has PduB knocked out and the ΔB::B-K102A-K207A strain has WT PduB replaced with a double mutant that disrupts PduB incorporation into the MCP shell. (B) Phase contrast and GFP fluorescence micrographs of cells expressing different GFP reporter constructs (scale bar = 1 µm). Different LT2 strains are indicated by the far left vertical axis label and the micrograph type is indicated by the near left vertical axis label. The reporter construct is indicated by the top horizontal labels. Representative bright, fluorescent puncta indicating MCPs are labelled with a blue arrow, polar bodies are labelled with a purple arrow, and diffuse fluorescence is indicated with a yellow arrow. (C) Quantification of puncta per cell for different fluorescently-tagged enzyme reporter constructs and various LT2 strains. The strain is indicated by the top, horizontal label and the encapsulation tag is indicated by the bottom horizontal label. Note that each reporter construct is readily encapsulated to varying degrees into the WT strain, whereas their encapsulation is significantly depleted in strains lacking PduB (p < 0.0001, one-tailed t-test). Puncta were counted from three biological replicates and >106 cells were counted per strain/construct.

When each fluorescent reporter is expressed in the WT strain they are localized to MCPs as indicated by the presence of bright fluorescent puncta (Figure 3B-C). This assay has been used as a readout of protein localization to MCPs and has been validated in numerous studies (27, 29, 30, 42). However, the relative encapsulation level of each enzyme set has not been quantitatively assessed using this assay before. Our results reveal that ssD-GFP and ssP-GFP are the most efficiently encapsulated, which is expected based on quantitative mass spectrometry analyses of purified MCPs (25). ssL-GFP is efficiently targeted to a relatively high number of MCPs, but puncta appear much dimmer qualitatively relative to ssD-GFP and ssP-GFP, as expected (Figure 3B). G-GFP and O-GFP are also encapsulated into MCPs, albeit less efficiently than any of the signal sequence tagged reporters. We hypothesize that this is due to their propensity to aggregate, as demonstrated by the formation of polar bodies in the ΔpocR strain (Figure 3B-C). Surprisingly, W-GFP is targeted to MCPs as well, in spite of the fact that it has been reported to be primarily cytosolic (Figure 3B-C) (25). It should be noted that it is only targeted to MCPs in a subset of cells, as indicated by the bimodal distribution in the MCP counts for this construct (Figure 3C). Together these results demonstrate that all of the enzymatic members of the Pdu metabolic pathway are likely targeted to some degree to MCPs, and that the GFP reporter constructs used here are adequate reporters of this targeting.

When the fluorescent reporter constructs are co-expressed with MCPs in the ΔB strain, only polar bodies are observed, similar to the ΔA ΔJ strain (Figure 3B-C). Quantitatively, significantly fewer puncta are observed per cell for each fluorescent reporter in both the ΔB and ΔA ΔJ strains compared with the WT strain. Significantly more puncta are present in the WT strain than in the ΔpocR strain as well, as expected (Figure 3C). This indicates that either MCP formation or enzyme encapsulation is significantly impacted by the loss of PduB.

### The PduB knockout strain forms assembled but empty MCPs

The encapsulation assay reveals that PduB loss dramatically alters the encapsulation state of all fluorescently tagged enzymes tested. However, this assay cannot distinguish between changes in encapsulation or changes in MCP formation. To test the hypothesis that PduB impacts encapsulation but not assembly, MCPs were purified from the ΔB strain using differential centrifugation (43, 44). When purified samples are analyzed by SDS-PAGE, the ΔB strain appears dramatically different from the standard MCP banding pattern apparent in the WT samples (Figure 4A). The ΔB samples more closely resemble the ΔA ΔJ samples, which are known to contain malformed MCP aggregates (Figure 4A-B) (27). An important distinction is that the ΔB sample has high apparent levels of PduA and PduJ (Figure 4A).

**Figure 4.**
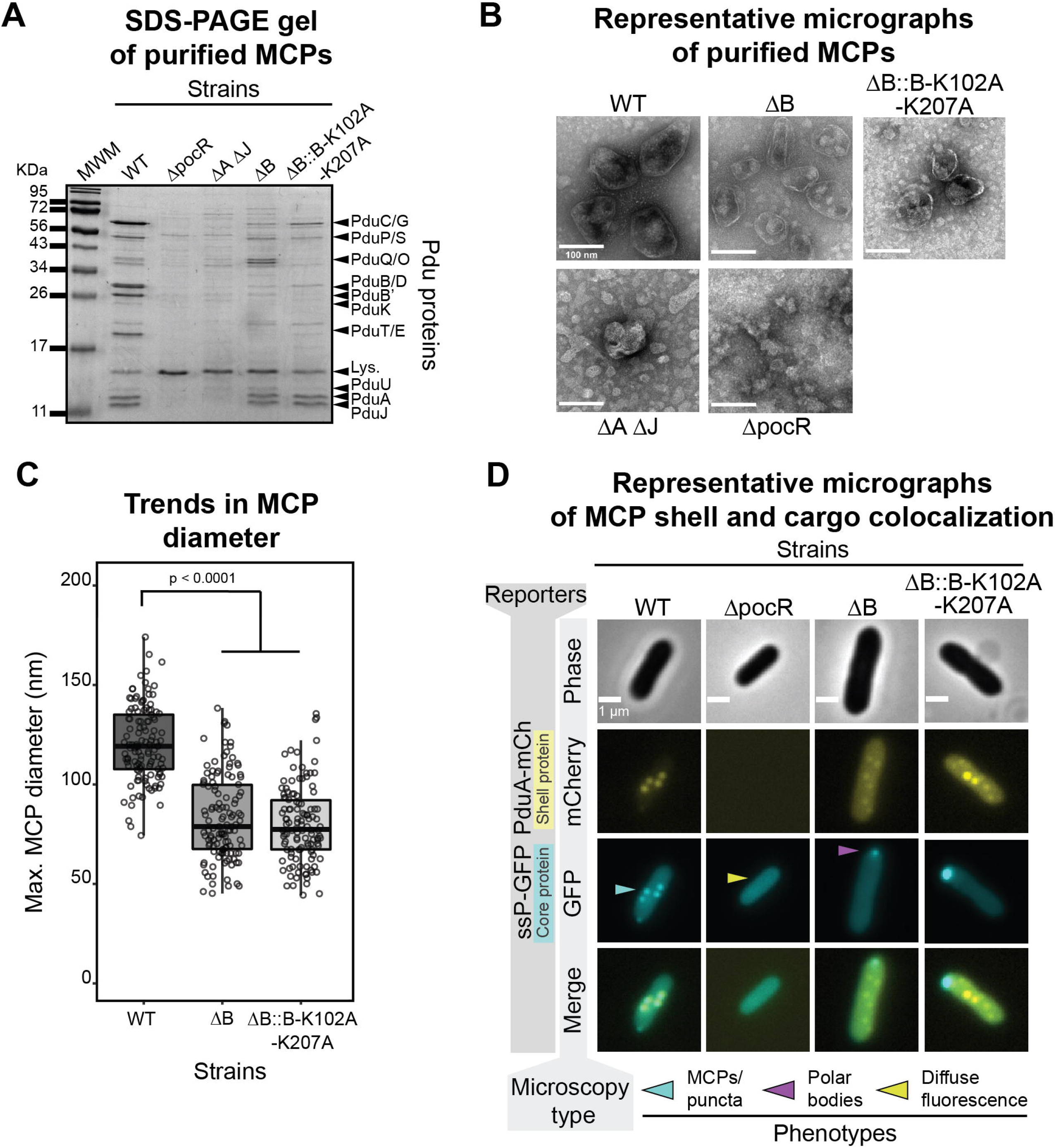
Loss of PduB decouples Pdu MCP shell assembly from the enzymatic core. (A) Coomassie-stained SDS-PAGE gel of MCP purifications from various LT2 strains (top horizontal label). Bands corresponding to various Pdu MCP components (right vertical labels) are present in the WT strain but absent in other strains. MWM = molecular weight marker. (B) Representative negative-stained transmission electron micrographs of Pdu MCP purifications from various LT2 strains. Well-formed MCP shells are irregular and polyhedral in shape and have a defined boundary and are present in the WT, ΔB, and ΔB::B-K102A-K207A samples. (C) Pdu MCPs purified from WT are significantly larger in diameter than shells purified from the ΔB or ΔB::B-K102A820 K207A strains (p < 0.0001, one-tailed t-test). Measurements were collected on three biological replicates and >120 MCPs were measured for each sample type. (D) Representative phase contrast and fluorescence micrographs for various strains of LT2 expressing ssP-GFP and PduA-mCherry reporter constructs (scale bar = 1 µm). The strain is labelled on the top horizontal axis and the microscopy type is labelled on the near left vertical axis. Bright, fluorescent puncta indicative of MCPs are labelled with a blue arrow, polar bodies are indicated with a purple arrow, and diffuse fluorescence is indicated with a yellow arrow. mCherry fluorescence is pseudocolored in yellow and GFP fluorescence is pseudocolored in cyan. Overlapping fluorescence is indicated in the merged image.

Purified samples were then analyzed by transmission electron microscopy (TEM) to determine if MCPs or partially formed aggregates were produced in the ΔB strain (Figure 4B). To our surprise, the ΔB sample shows many apparently well-formed MCPs. TEM micrographs from this sample contain structures with flat, distinguishable boundaries that appear roughly polyhedral in morphology (Figure 4B). Interestingly, the ΔB shells appear smaller than the WT shells (83 ± 21 nm vs 121 ± 19 nm, respectively (mean ± standard deviation)) and have qualitatively less electron density in their center, possibly indicating that they are empty (Figure 4B-C). The structures in the ΔB sample are clearly distinct from the aggregates observed in the ΔA ΔJ sample and the background structures present in the ΔpocR sample. These results, coupled with the SDS-PAGE and encapsulation assay results, indicate that the ΔB strain likely forms empty MCP shells as well as separate aggregates of Pdu enzymes.

To confirm this surprising finding, co-localization assays were performed in which the shell protein PduA was fused with a fluorescent mCherry reporter protein and expressed from the PduA locus in the WT, ΔB, and ΔpocR strain backgrounds. As expected, PduA-mCherry is localized to MCPs as indicated by the presence of bright fluorescent puncta (Figure 4D) (45). PduA is known to be important for shell formation and is one of the most abundant members of the Pdu MCP shell (25, 27, 46). When expressed in the WT background with the tagged GFP reporter constructs, GFP and mCherry colocalize, indicating reporter targeting to MCPs, as expected (Figure 4D). When expressed in the ΔpocR background, no mCherry puncta are present, as none of the proteins in the *pdu* operon, including PduA-mCherry, are expressed in this control (Figure 4D). In the ΔB strain background, mCherry puncta are still present, verifying the finding that MCP shells were still formed in the absence of PduB. As expected based on the results described in Figure 3, GFP reporter constructs are only found in polar bodies in the ΔB strain (Figure 4D). These results conclusively demonstrate that PduB deletion impacts protein encapsulation but not MCP shell formation, and that these are two, distinct processes that can be decoupled upon PduB deletion. Furthermore, this suggests that signal sequences are sufficient for targeting to the enzymatic core, challenging the idea that primary Pdu MCP signal sequence function is to bind to MCP shell proteins, enabling enzymatic core encapsulation (21, 23).

### PduB is incorporated into MCPs by a conserved hydrogen bonding mechanism

While PduB appears to be important for linking the MCP shell and enzymatic core, the mechanism by which PduB is incorporated into the shell (Figure 5A) is not known. A groundbreaking structural study by Sutter et al. showed that BMC-T^PD^ proteins similar to PduB appear to incorporate into the shell of a smaller, more regular, empty MCP from *Haliangium ochraceum* via a lysine-lysine hydrogen bonding motif (Figure 5B) (26, 47). In this motif, lysine side chains on adjacent BMC proteins (in this case, a BMC-H protein and BMC-T^PD^ protein) arrange antiparallel to each other and hydrogen bond with the backbone carbonyl oxygen of the adjacent lysine (Figure 5B) (26, 47). Interestingly, analogous lysine residues (Figure 5C) are also important for the self-assembly of BMC-H proteins PduA and PduJ, and are necessary for their ability to drive shell assembly (27). For these reasons, we hypothesized that analogous lysine residues at these positions in the BMC-T^P^ protein PduB (Figure 5C) are important for PduB incorporation into the MCP shell, bridging the gap between shell and enzymatic core.

**Figure 5.**
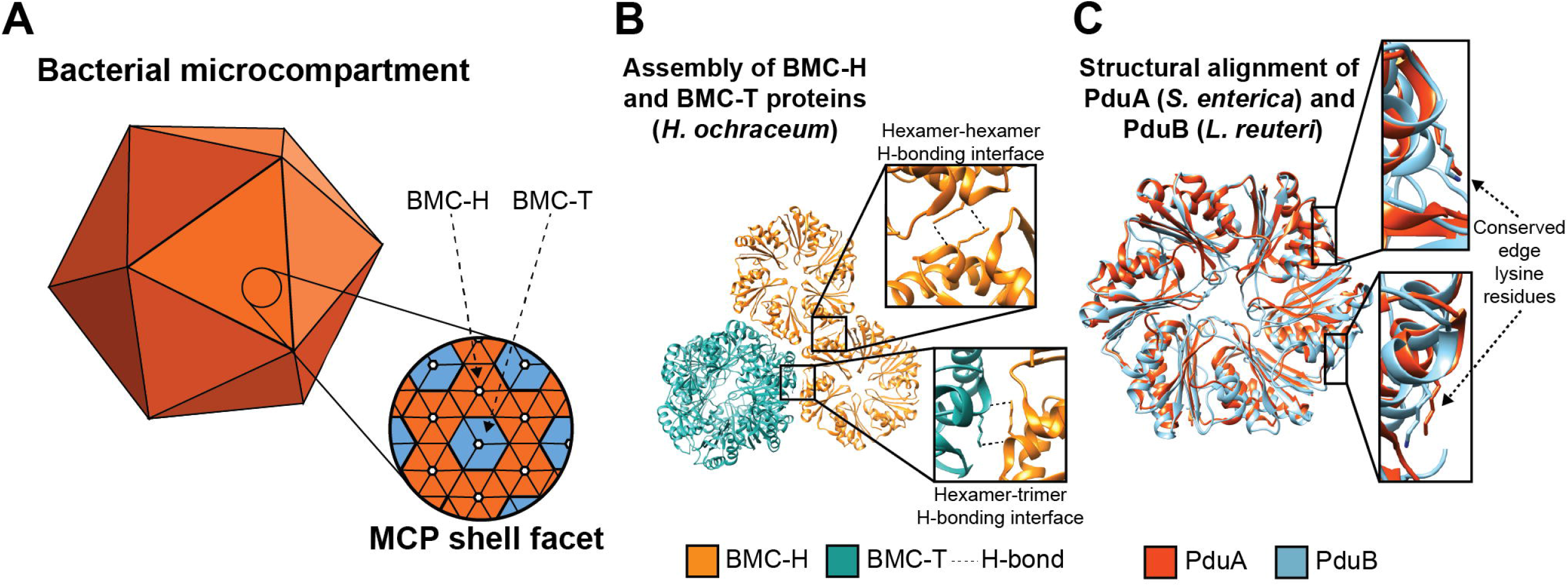
PduB is incorporated in the Pdu MCP shell via a conserved hydrogen bonding motif. (A) Schematic representation of BMC-H (orange) and BMC-T (blue) proteins tiling together to form the facets of the MCP shell. (B) Lysine residues at the hexamer-hexamer and hexamer-trimer interfaces are predicted to play a role in facet tiling in a MCP from *H. ochraceum* (PDB: 5V74) (26). BMC-H proteins are in orange and BMC-T proteins are in blue. (C) The edge lysine residue important for PduA (orange) self-assembly is also present in PduB (blue) (PduA PDB: 3NGK (55); PduB PDB: 4FAY (56)). All structural visualization was done using Chimera (57).

To test this hypothesis, a double mutant version of PduB was cloned where the lysine residues at position 102 and 207 were mutated to alanine residues. These residues appear to be well-conserved among PduB homologs (26), so we predicted that mutations to alanine should be disruptive. We expect that this protein is well-folded, as analogous mutations in other BMC proteins increase solubility and are used frequently in crystallographic studies (48, 49). The double mutant PduB (B-K102A-K207A) was then placed at the *pduB* locus in the *pdu* operon in place of WT *pduB*, generating the then placed at the *pduB* locus in the *pdu* operon in place of WT *pduB*, generating the ΔB::B-K102A-K207A strain (Figure 3A). The encapsulation assay, in which each fluorescent reporter is expressed simultaneously with MCPs, reveals that the ΔB::B K102A-K207A strain has a similar phenotype to both the ΔB and ΔA ΔJ strains, where polar bodies are primarily observed rather than the scattered puncta indicative of MCPs (Figure 3B-C). This indicates that the PduB double mutant is likely unable to incorporate into MCP shells, effectively recapitulating the ΔB phenotype. To confirm the conclusion based on the encapsulation assay, MCPs were purified from the ΔB::B-K102A-K207A strain. Samples from this strain appear similar to the ΔB strain by SDS-PAGE, where bands corresponding to enzymes are greatly reduced, but bands for PduA and PduJ are still visible (Figure 4A). Purified structures were then imaged by TEM, which reveals that apparently well-formed MCPs are present and similar in size and morphology to the MCPs purified from the ΔB strain (Figure 4B C). The PduB double mutant strain was also grown with PduA-mCherry expressed from the *pduA* locus, and this reveals bright puncta throughout the cytoplasm, similar to the ΔB strain, indicating the formation of MCP shells (Figure 4D). As in the ΔB strain, GFP tagged cargo appears only in polar bodies, again indicating well-formed shells decoupled from the enzymatic cargo (Figure 4D). Together, these results strongly support the hypothesis that PduB is incorporated into the MCP shell by a similar mechanism to other BMC proteins, and that disruption of the conserved lysine hydrogen bonding network disrupts this interaction.

### Loss of PduB impacts LT2 growth on 1,2-propanediol as a carbon source

Having established that PduB is essential for linking the core to the shell of the Pdu MCP, we next explored how decoupling of the core and shell impacts the function of the native Pdu pathway. To this end, we compared growth and metabolite profiles in culture conditions that differentiate strains with well-formed MCPs and broken MCPs. Specifically, these growth conditions encourage buildup of the toxic propionaldehyde intermediate, leading to a growth lag in strains with broken MCPs that cannot sequester this intermediate away from the cytosol (9, 16, 27). Under these growth conditions, we hypothesized that our strains with decoupled core and shell assembly would exhibit a growth phenotype similar to that of a strain expressing broken Pdu MCPs (ΔA ΔJ).

We tested this hypothesis by comparing the growth and metabolite profiles of three control strains (WT, ΔpocR, ΔA ΔJ) to two experimental strains (ΔB, ΔB::B K102A-K207A). Our three control strains were the same as those used in the encapsulation assay—WT, which harbors functional MCPs, ΔpocR, which cannot express the *pdu* operon, and ΔA ΔJ, which forms proto-MCP aggregates but no shell and thus serves as our broken MCP control. Our two experimental strains (ΔB, ΔB::B K102A-K207A), as described above, both have decoupled MCP core and shell assembly. Our control strains behave as expected. The ΔpocR strain, which is unable to express any Pdu pathway enzymes, does not grow nor does it consume 1,2-PD (Figure 6A, B). Our broken MCP control strain (ΔA ΔJ) initially grows and consumes 1,2-PD faster than the MCP-containing strain (WT), as expected—the absence of a MCP shell accelerates apparent pathway kinetics by providing more direct access to 1,2-PD (Figure 6A, B). However, propionaldehyde build up in the ΔA ΔJ strain eventually leads to a lag in growth from 12-30 hours (Figure 6A, B). During this timeframe, the WT strain begins to outgrow the ΔA ΔJ strain, as the MCPs in the WT strain successfully sequester the toxic propionaldehyde intermediate away from the cytosol (Figure 6A, B).

**Figure 6.**
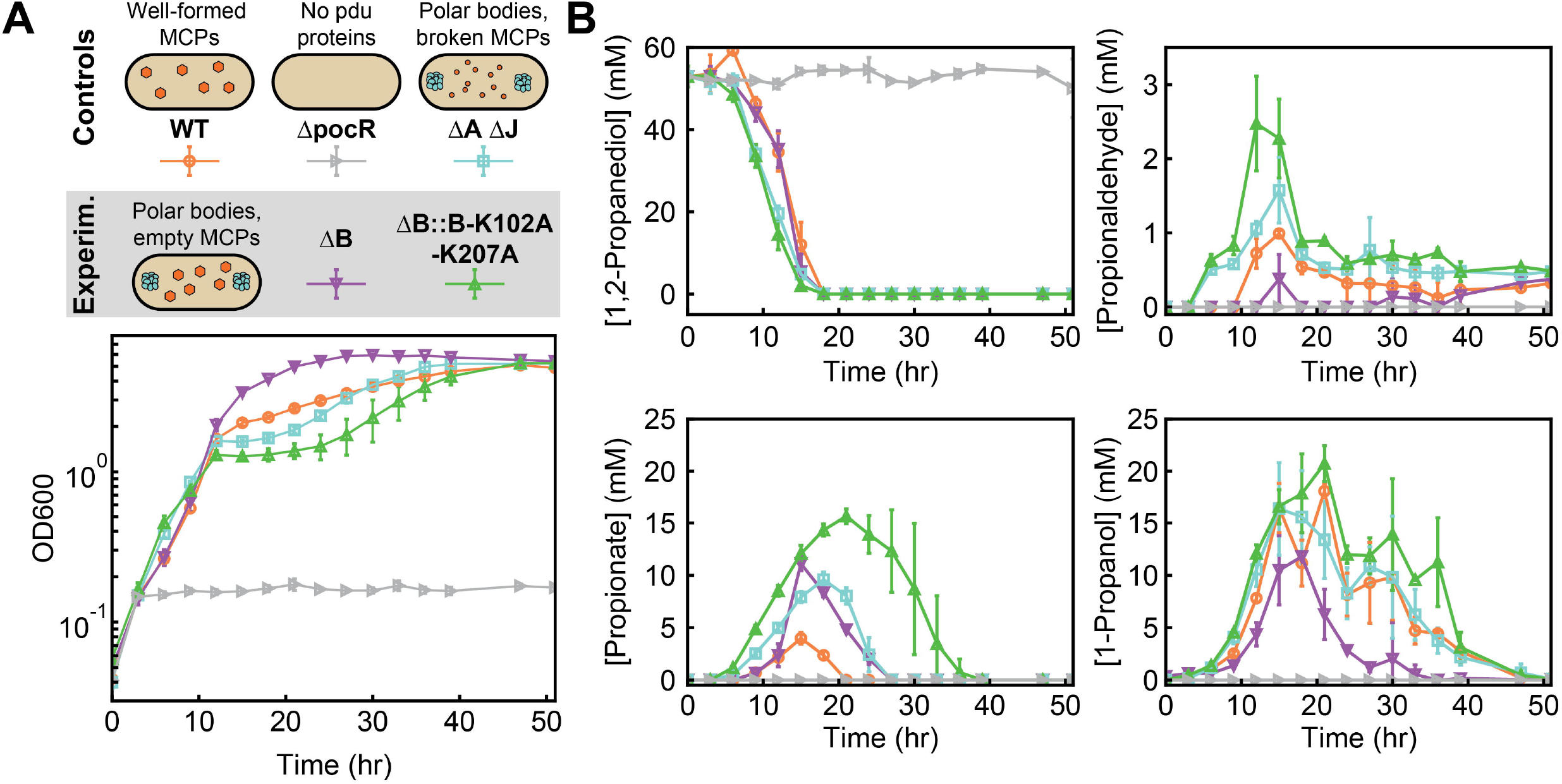
PduB alterations change LT2 growth and metabolite profiles. (A) LT2 control and experimental strains containing different MCP morphologies (WT MCPs, polar bodies (ΔA ΔJ, ΔB, and ΔB::B-K102A-K207A), no protein expression (ΔpocR)). These differing morphologies lead to various LT2 growth profiles, as measured by the optical density at 600 nm (OD600) at different time points. (B) Metabolite profiles of different LT2 strains during growth on 1,2-propanediol as a sole carbon source with excess adenosylcobalamin (150 nM). The concentration (mM) of four key metabolites (1,2-propanediol, propionaldehyde, propionate, and 1-propanol) was measured over the duration of the growth curve visualized in Fig. 6A.

Surprisingly, we find that our two experimental strains (ΔB, ΔB::B-K102A K207A), both of which have decoupled MCP core and shell assembly, exhibit distinct growth and metabolite profiles. The growth and metabolite profiles of the PduB double mutant strain (ΔB::B-K102A-K207A) are similar to those of the broken MCP control (ΔA ΔJ), in agreement with our initial hypothesis. This strain (ΔB::B-K102A-K207A) grows rapidly at first, and then suffers a growth lag after propionaldehyde accumulates after 9 hours (Figure 6A, B). Interestingly, the growth lag is longer (9-40 hours) and the peak propionaldehyde level is higher (2.5 ± 0.6 mM) in the ΔB::B-K102A-K207A strain than in the ΔA ΔJ strain. More perplexing is the behavior of the ΔB strain—not only does this strain exhibit minimal propionaldehyde buildup, it also eventually outgrows the WT strain (Figure 6A, B). Correspondingly, the downstream metabolites propionate and 1- propanol are consumed more rapidly in the ΔB strain than in other strains (Figure 6A, B). Given that ΔB and ΔB::B-K102A-K207A strains exhibit essentially identical behavior with respect to assembly phenotype (shown above), we turned to kinetic modeling to generate hypotheses that could explain this observed discrepancy in growth behavior and differences in propionaldehyde accumulation.

Using a systems-level kinetic model of the Pdu pathway modified from previous work (8) (see Materials and Methods), we examined which features of the pathway or polar body geometry had the strongest impact on propionaldehyde buildup in cells containing polar bodies. We considered MCPs as a control case, modeling these MCPs as spheres with a diffusive barrier at their surface that limits metabolite transport to the enzymatic core. Polar bodies, then, are modeled as one large sphere with free diffusion of metabolites at the surface (Figure 7A). The volume of this polar body sphere was set to equal the total volume of MCPs in the cell (*i.e.* polar body volume = number of MCPs per cell x volume of a single MCP). We used this polar body model to assess the sensitivity of maximum propionaldehyde buildup to parameters affecting kinetics, geometry and transport. Comparison of the MCP (Supplementary Figure S1) and polar body (Supplementary Figure S2) models reveals that the maximum propionaldehyde buildup in the polar body case is much more sensitive to the maximum velocity (*V*_max_) of the PduCDE enzyme than in the MCP case. As the concentration of PduCDE in the polar body is decreased (leading to an equivalent fractional decrease in *V*_max_), the predicted propionaldehyde profile for the polar body approaches that of the MCP (Figure 7A). Indeed, decreasing the PduCDE concentration in the polar bodies decreases the maximum propionaldehyde concentration outside the cell (Figure 7B).

**Figure 7.**
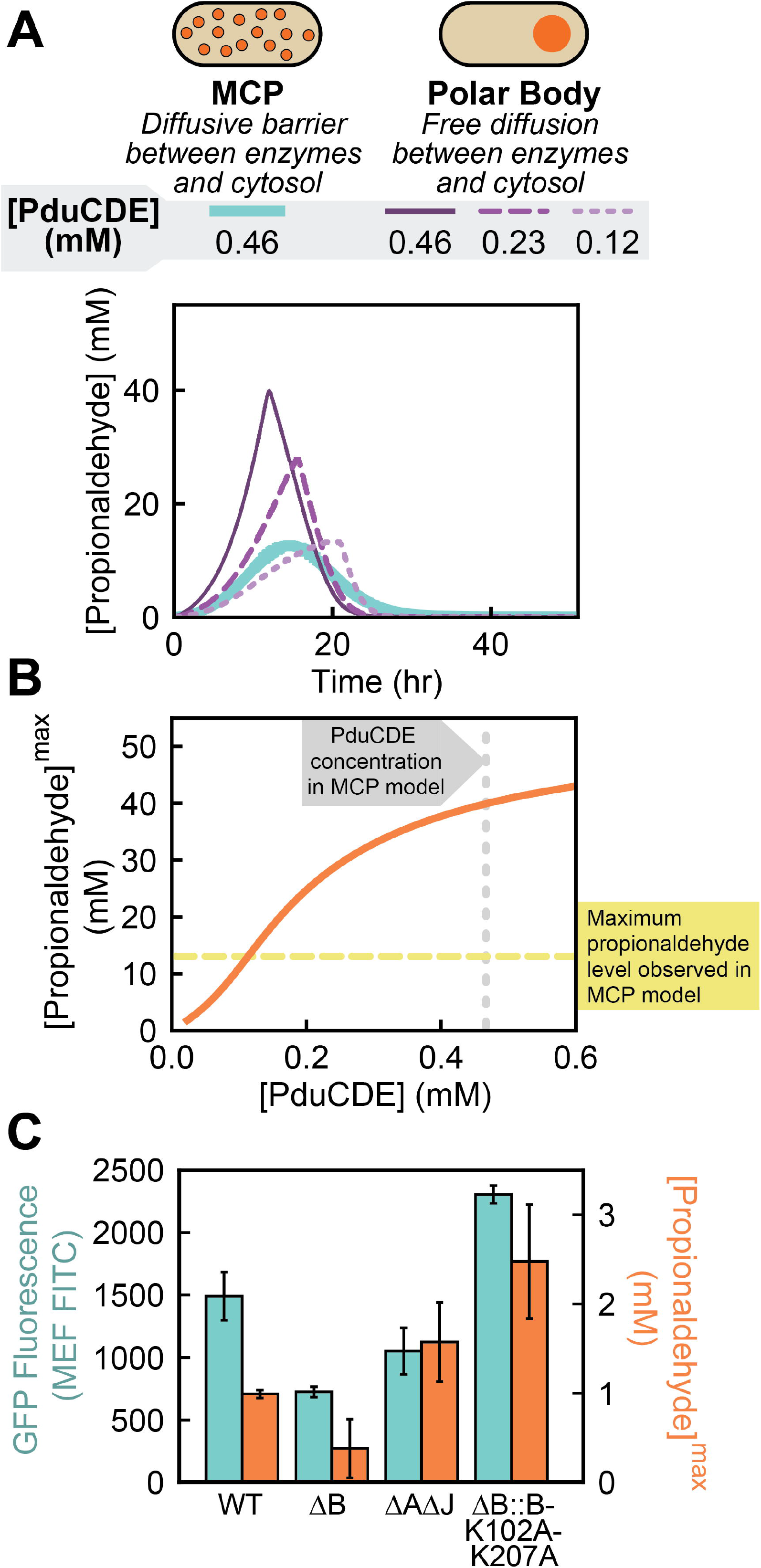
System-level kinetic modelling reveals how a polar effect alters growth and metabolite profiles. (A) Two organizational strategies for Pdu enzymes (MCPs and polar bodies) were tested in a systems-level kinetic model. Plotted here are propionaldehyde concentrations in the external media. PduCDE enzyme concentration was also varied in the polar body model case to account for the potential genetic polar effects that alter gene expression. (B) The kinetic model demonstrates that changes to PduCDE concentration (mM) impact the maximum propionaldehyde concentration (mM) observed in the external media when pathway enzymes are contained in a polar body.(C) Comparison of GFP fluorescence (MEF FITC) measured by flow cytometry as a proxy for gene expression vs peak propionaldehyde concentration (mM) reveals that PduCDE expression is likely reduced due to a genetic polar effect in the ΔB strain.

Specifically, we find that the maximum propionaldehyde concentration observed in the polar body model matches that of the MCP model if the PduCDE concentration is decreased by 76%, suggesting that variation in PduCDE concentration between strains can impact propionaldehyde buildup as strongly as Pdu pathway encapsulation in MCPs (Figure 7B). We thus hypothesized that the difference between our strains containing polar bodies was the level of PduCDE expression. We hypothesized that this was the result of polar effects in which different alterations to the *pduB* locus differentially impact the expression of downstream *pduCDE* genes.

We tested our hypothesis that PduCDE expression differed across strains containing polar bodies by measuring GFP expression at the *pduD* locus. Specifically, we replaced the *pduD* gene at the *pduD* locus with a gene encoding for ssD-GFP in our five growth strains (WT, ΔpocR, ΔA ΔJ, ΔB, ΔB::B-K102A-K207A) and used flow cytometry to quantify GFP expression. We find that the ΔB strain has the lowest expression at the *pduD* locus, with 49 ± 7% lower GFP fluorescence than the WT strain (*p* < 0.05). By contrast, the ΔB::B-K102A-K207A strain had higher apparent GFP expression at the *pduD* locus than the WT strain (*p* < 0.05). Finally, we find that our broken MCP control strain, ΔA ΔJ, has just slightly lower expression at the *pduD* locus than WT (*p* < 0.05) (Figure 7C). Importantly, we note that increased GFP expression at the *pduD* locus correlates with peak propionaldehyde level in the strains with unencapsulated enzymes (Pearson’s correlation coefficient = 0.92), supporting our hypothesis that PduCDE expression causes differences in propionaldehyde buildup in these strains.

Together, these results highlight two important considerations for future work in the MCP field. First, modeling is a powerful tool for generating hypotheses and exploring parameter space, enabling rapid identification of the key governing parameters for a given subcellular organizational strategy. Second, polar effects can impact downstream enzyme expression, which, in turn, can dramatically alter observed pathway performance. Thus, quantifying these effects and minimizing them whenever possible is essential when conducting genetic studies of MCP function.

## Discussion

Metabolosomes are bacterial organelles confirmed to exist in a number of enteric bacteria, where their expression and activation has been linked to pathogenesis (10). Of the metabolosomes investigated to-date, the Pdu MCP is the best studied, and has served as a model system for understanding the basics of metabolosome assembly and metabolic function. However, even in this well-studied model system, a large body of conflicting evidence exists regarding the mechanism by which enzymatic cargo is loaded into the MCP shell (20–24).

Here, we used a comprehensive suite of fluorescent reporters to show that the shell protein PduB is responsible for linking the enzymatic core to the MCP shell—when PduB is absent or unable to incorporate into the shell, no Pdu enzymes are loaded into the MCP. This interaction between PduB and the enzymatic core likely happens via the N-terminus of PduB, as previously suggested in Lehman et al., since deletion of this unique structural domain reduces enzyme loading into MCPs (20). Instead, enzymes localize to proto-MCP aggregates, termed polar bodies, in the absence of PduB. Importantly, we find that N-terminal signal sequence tags such as those identified for PduD and PduP are sufficient for loading cargo to these proto-MCP aggregates (29, 30). This is significant as these signal sequence tags were previously thought to exist primarily to link the shell and core of the MCP. We also report that although PduB is required for cargo loading, it is not necessary for MCP shell assembly, as empty MCP structures form in the absence of PduB. The fact that empty MCP shells and proto-MCP aggregates form separately in the absence of PduB suggests that MCP shell and core assembly are independent processes in the Pdu MCP system. Combined with previous work demonstrating that a self-assembling proteins PduA or PduJ are required for Pdu MCP formation (27), this discovery brings us one step closer to realizing the components necessary to construct a minimal Pdu MCP. Specifically, two of the requisite components are: (1) a hexameric protein with a strong propensity for self-assembly (PduA or PduJ) and (2) a protein that links the core and shell by interacting with both the enzymatic core and the self-assembling hexameric proteins via a conserved hydrogen bonding mechanism. We expect that computational models of this assembly process will provide insight into the specific roles that these different proteins play in MCP assembly (46, 50).

We also investigated how LT2 growth on 1,2-propanediol was impacted by the decoupling of MCP core and shell assembly. As expected, when enzymes are localized to polar bodies, buildup of the toxic aldehyde intermediate is not well-controlled as it is in MCPs. However, using a combination of systems-level kinetic modeling and experiments, we also discovered that strains that contain polar bodies rather than MCPs exhibit dramatically different growth profiles depending on PduCDE expression level. In our strains, we believe the differences in PduCDE expression we observed are a consequence of polar effects on gene expression, as the genes encoding for *pduCDE* are immediately downstream of the *pduB* gene. Specifically, we find that while our *pduB* knockout strain (ΔB) and our *pduB* double mutant strain (ΔB::B-K102A-K207A) both result in a similar decoupling of MCP core and shell assembly, the strain with the more dramatic genetic alteration at the *pduB* locus (ΔB, the knockout) decreases expression at the *pduD* locus substantially, whereas the strain with minimal genetic alteration at the *pduB* locus (ΔB::B-K102A-K207A) does not. This result emphasizes an important point of concern for studies in this field—namely that polar effects on downstream gene expression can confound interpretation of experiment results, and thus must be considered. Indeed, it has been shown that alterations to the *pduL* locus, which encodes for enzymatic cargo, can disrupt Pdu MCP shell assembly, likely by modifying expression levels of downstream shell proteins like PduN (42). Conversely, we show here that modifications to the *pduB* locus, which encodes for a shell protein, can alter enzyme concentrations in the core, altering Pdu pathway performance. Thus, the identification of point mutants that minimally disrupt operon sequence, but prevent specific protein functions, will be essential in future MCP studies.

## Materials and Methods

### Plasmid and strain creation

All plasmids used for this study were created using the Golden Gate cloning (51) method into a Golden Gate-compatible pBAD33t parent vector (chloramphenicol resistance, p15a origin of replication). All strains, plasmids, and primers are listed and described in Supplemental Tables S1-3, respectively. For constructs in which a signal sequence was appended to the N-terminus of GFPmut2, a BsaI cut site and the DNA sequence encoding the signal sequence were added to GFP using PCR (see Table S3 for primer sequence information). For strains in which GFPmut2 was appended to the C-terminus of a full enzyme, a two-piece Golden Gate reaction was carried out. This is true for all GFP-enzyme fusions with the exception of PduG-GFP, which was cloned using SacI and XbaI restriction sites. First, forward and reverse primers were used to amplify the enzyme and GFPmut2 open reading frames. Each primer encoded compatible sticky ends to enable ligation into the pBAD33t parent vector in the proper orientation. A glycine-serine (GS) linker was also encoded between the enzyme and GFPmut2 open reading frames. Golden Gate reactions were carried out using FastDigest Eco31I (Thermo Fisher Scientific) and T4 DNA ligase (New England Biolabs), transformed into *E. coli* DH10b cells, purified using Zyppy^TM^ Plasmid Miniprep kit (Zymo Research), and sequence verified by Sanger sequencing (Genewiz).

All modified LT2 strains used in this study are listed in Table S1 and were generated using λ red recombineering as described in previous reports (27, 52). Briefly, a marker encoding dual selection (*cat/sacB*) was PCR amplified containing flanking homologous overhangs matching the target gene locus and inserted into the *pdu* operon. The *cat/sacB* selectable marker was then either replaced with a gene of interest or knocked out, leaving only the C-terminus of the target gene in order to reduce polar effects on downstream gene expression. For the PduB-K102A-K207A double mutant strain, QuikChange was used (with KOD Hot Start DNA Polymerase (Sigma-Aldrich)) to perform site directed mutagenesis on the two lysine residues on a plasmid backbone first before it was amplified and inserted into the *pdu* operon. Modified strains were sequence confirmed using Sanger sequencing (Genewiz).

### Fluorescence and phase contrast microscopy

Fluorescence and phase contrast micrographs were collected on Nikon Eclipse Ni-U upright microscope, 100X oil immersion objective, and an Andor Clara digital camera with NIS Elements Software (Nikon). GFP fluorescence micrographs were collected using a C-FL Endow GFP HYQ bandpass filter and mCherry fluorescence micrographs were collected using a C-FL Y-2E/C filter. Exposure times of 100 ms were used for ssD-GFP and ssP-GFP, 300 ms for ssL-GFP, 400 ms for G-GFP, O-GFP, and W-GFP, and 500 ms for mCherry. Cell culture samples were placed onto Fisherbrand^TM^ frosted microscope slides and covered using 22 mm x 22 mm, #1.5 thickness cover slips. All images were equally adjusted within experiments and sample type for brightness and contrast using ImageJ (53). Puncta counts were collected on brightness and contrast adjusted images.

### Microcompartment expression and purification

MCP expression and purification was done as described in detail previously (43). Briefly, overnight starter cultures were started from single colonies streaked out onto Lysogeny Broth, Miller (LB-M, Thermo Fisher Scientific) agar plates. These 5 mL LB-M µg/mL chloramphenicol added when appropriate) were grown in 24-well blocks for 15-18 hours at 37°C and shaking at 225 RPM. Saturated starter cultures were used to subculture expression cultures. If cultures were used for microscopy experiments, cultures were subcultured 1:500 in 5 mL LB-M supplemented with 0.02% (w/v) final concentration of L-(+)-arabinose to induce fluorescent reporter expression, 0.4% (v/v) final concentration 1,2-propanediol to induce MCP expression, and 34 µg/mL chloramphenicol. Cultures were then grown for a minimum of 6 hours at 37°C, 225 RPM before imaging. For cultures used for MCP purification, saturated overnight cultures were diluted 1:1000 into 200 mL No Carbon Essential (NCE) media supplemented with 50 µM ferric citrate, 42 mM succinate as a carbon source, 1 mM magnesium sulfate, and 55 mM 1,2-propanediol to induce MCP expression. The cultures were grown in 1 L glass Erlenmeyer flasks at 37°C, 225 RPM until cultures reached an OD_600_ of at least 1. MCPs were purified using an established differential centrifugation method described previously (43). Briefly, cells were pelleted at 5,000 x g and lysed chemically using an octylthioglucoside (OTG) solution. Cell lysate was clarified at 12,000 x g (4°C, 5 minutes) and MCPs were pelleted at 21,000 x g (4°C, 20 minutes). Purified MCPs were stored in a buffered saline solution containing 50 mM Tris (pH 8.0), 50 mM potassium chloride, 5 mM magnesium chloride, and 1% (v/v) 1,2-propanediol at 4°C. Cultures used for flow cytometry were subcultured from overnights 1:1000 into 5 mL of the same NCE media used for compartment purifications and grown for 16 hours at 37 °C at 225 RPM prior to analysis.

### Transmission electron microscopy

Transmission electron microscopy (TEM) on purified MCPs was done as described in detail in a prior study (15). Samples containing purified MCPs were applied and fixed onto 400 mesh Formvar-coated copper grids (EMS Cat# FF400-Cu). Fixation was done using 2% (v/v) glutaraldehyde and staining was done using 1% (w/v) uranyl acetate. Samples were then imaged using a JEOL 1230 transmission electron microscope with a Gatan 831 bottom-mounted CCD camera. MCP diameter measurements were collected using the Ferret diameter tool in ImageJ (53).

### SDS-PAGE

For analysis of purified MCPs using SDS-PAGE, samples were first normalized by protein concentration (measured by bicinchoninic acid assay) before loading onto the 15 wt% polyacrylamide Tris-glycine minigels. SDS-PAGE was then run a second time with sample loading normalized by densitometry on the PduA band to correct for potential contaminants within the samples. Samples were boiled at 95°C for 5 minutes in Laemmli buffer and then immediately run at 120 V for 90 minutes. Gels were stained with Coomassie Brilliant Blue R-250 and imaged using the Bio-Rad ChemiDoc XRS + System.

### Growth assays and metabolite analysis

Overnight starter cultures of LT2 strains of interest were started from a single colony and grown in test tubes in 5 mL of Terrific Broth (Dot Scientific, Inc.) without glycerol. Overnights were grown 15-16 h at 37 °C at 225 RPM with orbital shaking. These overnights were subcultured into 50 mL No Carbon Essential (NCE) media supplemented with 50 µM ferric citrate, 1 mM magnesium sulfate, 150 nM adenosylcobalamin (Santa Cruz Biotechnology), and 55 mM 1,2-propanediol to an OD_600_ of 0.05. Cultures were grown in 250 mL, unbaffled Erlenmeyer flasks at 37 °C, 225 rpm. At each time point, 500 µL of cell culture was taken for OD_600_ measurement and HPLC analysis. OD_600_ measurements were taken using a BioTek Synergy HTX multi-mode plate reader. After 9 hours, cultures from all strains except the ΔpocR strain were diluted 1:5 in fresh NCE to ensure measurements were within the linear range of the instrument. Error bars on growth curves represent standard deviation over three biological replicates. The remainder of the cell culture sample (that not used for OD_600_ measurement) was centrifuged at 13,000 x g for 5 minutes to pellet cells. The supernatant was collected and frozen at -20 °C. After completion of the growth experiment, supernatant samples were thawed and filtered (Corning™ Costar™ Spin-X LC filters) in preparation for HPLC analysis. Samples were run on an Agilent 1260 HPLC system. Metabolites were separated using a Rezex™ ROA-Organic Acid H+ (8%) LC Column (Phenomenex) at 35 °C, in 5 mM sulfuric acid flowing at 0.4 mL/min. Metabolites were detected with a refractive index detector (RID) as previously described (9). Peak areas were calculated using the Agilent ChemLab software and converted to metabolite concentrations using standards of the metabolites of interest (1,2- propanediol, propionaldehyde, propionate, 1-propanol) at 200 mM, 100 mM, 50 mM, 20 mM, and 5 mM. Error bars on metabolite concentrations represent standard deviation over three biological replicates.

### Modelling

The kinetic model used here is modified from previous work, which described MCP function in the cell using a reaction-diffusion framework (8). Modifications include an assumption that the cytosol is well-mixed to produce a compartmental model and the incorporation of the effects of cell growth. We analyze how metabolite profiles evolve over time, while previous studies focused on steady state. A detailed description of the model used in this work is provided here and in Supplemental Document 1.

We model cells as capsule-shaped (cylinders with a hemisphere at either end) and allow passive transport of 1,2-propanediol, propionaldehyde, propionyl-CoA, 1- propanol, and propionate at the cell surface, specified by the permeabilities included in Supplemental Table S4. MCPs are modeled as spheres 140 nm in diameter (15 per cell) that have a specified permeability at the surface; this permeability is assumed to be the same for all metabolites. Polar bodies are modeled as a single sphere per cell 345 nm in diameter (selected such that the volume of the polar body is equal to that of 15 MCPs), and allow free diffusion of all metabolites at their surface. Recognizing that diffusion in the MCP, cytosol, and media are much faster than the transport processes and enzyme reaction rates, we assume that the concentration is uniform in each of these locations.

We assume that conversion of 1,2-propanediol to propionaldehyde by PduCDE is irreversible. Similarly, we assume that conversion of propionyl-CoA to propionate by PduL and PduW occurs in a single step, and that this reaction is also irreversible. Reverse reactions are included for conversion of propionaldehyde to 1-propanol and propionyl-CoA by PduQ and PduP respectively. Reactions catalyzed by PduCDE, PduP, and PduQ are assumed to occur only in the MCP or polar body volume, whereas the reaction catalyzed by PduL/PduW is assumed to occur only in the cytosol. All enzymes are assumed to have Michaelis-Menten kinetics.

Cell concentration and growth are calculated using the data from the growth curves. The model was built and implemented in Python (54). All codes used are available on GitHub at https://github.com/cemills/MCP-vs-PolarBody. A detailed description of the equations used in the model are available in Supplemental Document 1. The parameters used for calculations presented here are available in Supplemental Table S4. Metabolite profiles over time of 1,2-propanediol, propionaldehyde, propionate, 1-propanol for the PduCDE concentrations shown in Figure 7A can be found in Supplementary Figure S3.

### Flow cytometry

Prior to measurement, cells were diluted to OD_600_ = 0.03 into 200 µL phosphate-buffered saline (PBS) supplemented with 2 mg/mL kanamycin to stop translation. Samples were prepared in a U-bottomed 96-well plate and kept away from light until measurement. 10,000 events were collected per sample on an Attune NxT acoustic focusing cytometer paired with an Attune NxT auto sampler. Cell populations were gated using forward and side scatter channels, and average reported fluorescence was calculated using the geometric mean of the population. Values were converted to molecules of equivalent fluorescence (MEF) using BD Sphero™ Rainbow Calibration Particles (Fisher Scientific catalog number BDB559123) according to manufacturer’s instructions. Analysis was performed using FlowJo software (www.FlowJo.com).

## Acknowledgements

The authors would like to thank and acknowledge members of the Tullman-Ercek and Mangan groups for insightful advice and discussion related to designing experiments associated with this study as well as preparing the manuscript. The authors would like to specifically acknowledge Dr. Taylor M. Nichols, Dr. Eddy Kim, Dr. Chris Jakobson, and Mike Vincent for strains, plasmids, and primers used in this study. This work was funded in part by the Army Research Office (grant W911NF-19-1-0298 to DTE), and the Department of Energy (grant DE-SC0019337 to DTE and NMM). NWK was funded by the National Science Foundation Graduate Research Fellowship Program (grant DGE- 1842165), and by the National Institutes of Health Training Grant (T32GM008449) via the Northwestern University Biotechnology Training Program.

CRediT Contributor Roles: **NWK**: Conceptualization, formal analysis, funding acquisition, investigation, methodology, project administration, validation, visualization, writing – original draft, writing – review & editing. **CEM**: Conceptualization, formal analysis, investigation, methodology, project administration, software, validation, visualization, writing – original draft, writing – review & editing. **CHA**: Conceptualization, investigation, methodology, writing – review & editing. **AA**: Conceptualization, formal analysis, software, writing – review and editing. **MCJ**: Funding acquisition, resources, supervision, writing – review & editing. **NMM**: Conceptualization, funding acquisition, resources, supervision, writing – review & editing. **DTE**: Conceptualization, funding acquisition, resources, supervision, writing – review & editing.

## References

1. Gabaldón T, Pittis AA. 2015. Origin and evolution of metabolic sub-cellular compartmentalization in eukaryotes. Biochimie 119:262–268.

2. Uebe R, Schüler D. 2016. Magnetosome biogenesis in magnetotactic bacteria. Nature Reviews Microbiology 14:621–637.

3. Greening C, Lithgow T. 2020. Formation and function of bacterial organelles. Nat Rev Microbiol 18:677–689.

4. Kennedy NW, Mills CE, Nichols TM, Abrahamson CH, Tullman-Ercek D. 2021. Bacterial microcompartments: tiny organelles with big potential. Current Opinion in Microbiology 63:36–42.

5. Kerfeld CA, Melnicki MR. 2016. Assembly, function and evolution of cyanobacterial carboxysomes. Current Opinion in Plant Biology 31:66–75.

6. Kerfeld CA, Aussignargues C, Zarzycki J, Cai F, Sutter M. 2018. Bacterial microcompartments. Nature Reviews Microbiology 16:277–290.

7. Asija K, Sutter M, Kerfeld CA. 2021. A Survey of Bacterial Microcompartment Distribution in the Human Microbiome. Frontiers in Microbiology 12:1090.

8. Jakobson CM, Tullman-Ercek D, Slininger MF, Mangan NM. 2017. A systems-level model reveals that 1, 2-Propanediol utilization microcompartments enhance pathway flux through intermediate sequestration. PLoS computational biology 13:e1005525.

9. Sampson EM, Bobik TA. 2008. Microcompartments for B12-Dependent 1,2- Propanediol Degradation Provide Protection from DNA and Cellular Damage by a Reactive Metabolic Intermediate. Journal of Bacteriology 190:2966–2971.

10. Prentice MB. 2021. Bacterial microcompartments and their role in pathogenicity. Current Opinion in Microbiology 63:19–28.

11. Stewart KL, Stewart AM, Bobik TA. 2020. Prokaryotic Organelles: Bacterial Microcompartments in E. coli and Salmonella. EcoSal Plus 9.

12. Axen SD, Erbilgin O, Kerfeld CA. 2014. A Taxonomy of Bacterial Microcompartment Loci Constructed by a Novel Scoring Method. PLoS Computational Biology 10:e1003898.

13. Sutter M, Melnicki MR, Schulz F, Woyke T, Kerfeld CA. 2021. A catalog of the diversity and ubiquity of bacterial microcompartments. Nat Commun 12:3809.

14. McFarland AG, Kennedy NW, Mills CE, Tullman-Ercek D, Huttenhower C, Hartmann EM. 2021. Density-based binning of gene clusters to infer function or evolutionary history using GeneGrouper. bioRxiv 2021.05.27.446007.

15. Kennedy NW, Hershewe JM, Nichols TM, Roth EW, Wilke CD, Mills CE, Jewett MC, Tullman-Ercek D. 2020. Apparent size and morphology of bacterial microcompartments varies with technique. PLOS ONE 15:e0226395.

16. Slininger Lee MF, Jakobson CM, Tullman-Ercek D. 2017. Evidence for Improved Encapsulated Pathway Behavior in a Bacterial Microcompartment through Shell Protein Engineering. ACS Synth Biol 6:1880–1891.

17. Kerfeld CA, Sawaya MR, Tanaka S, Nguyen CV, Phillips M, Beeby M, Yeates TO. 2005. Protein structures forming the shell of primitive bacterial organelles. Science 309:936–938.

18. Wheatley NM, Gidaniyan SD, Liu Y, Cascio D, Yeates TO. 2013. Bacterial microcompartment shells of diverse functional types possess pentameric vertex proteins. Protein Science 22:660–665.

19. Bobik TA, Havemann GD, Busch RJ, Williams DS, Aldrich HC. 1999. The Propanediol Utilization (pdu) Operon ofSalmonella enterica Serovar Typhimurium LT2 Includes Genes Necessary for Formation of Polyhedral Organelles Involved in Coenzyme B12-Dependent 1, 2-Propanediol Degradation. Journal of bacteriology 181:5967–5975.

20. Lehman BP, Chowdhury C, Bobik TA. 2017. The N Terminus of the PduB Protein Binds the Protein Shell of the Pdu Microcompartment to Its Enzymatic Core. Journal of Bacteriology 199:e00785–16.

21. Lawrence AD, Frank S, Newnham S, Lee MJ, Brown IR, Xue W-F, Rowe ML, Mulvihill DP, Prentice MB, Howard MJ, Warren MJ. 2014. Solution Structure of a Bacterial Microcompartment Targeting Peptide and Its Application in the Construction of an Ethanol Bioreactor. ACS Synthetic Biology 3:454–465.

22. Parsons JB, Lawrence AD, McLean KJ, Munro AW, Rigby SEJ, Warren MJ. 2010. Characterisation of PduS, the pdu Metabolosome Corrin Reductase, and Evidence of Substructural Organisation within the Bacterial Microcompartment. PloS one 5:e14009.

23. Fan C, Cheng S, Sinha S, Bobik TA. 2012. Interactions between the termini of lumen enzymes and shell proteins mediate enzyme encapsulation into bacterial microcompartments. Proceedings of the National Academy of Sciences 109:14995– 15000.

24. Jorda J, Liu Y, Bobik TA, Yeates TO. 2015. Exploring Bacterial Organelle Interactomes: A Model of the Protein-Protein Interaction Network in the Pdu Microcompartment. PLOS Computational Biology 11:e1004067.

25. Yang M, Simpson DM, Wenner N, Brownridge P, Harman VM, Hinton JCD, Beynon RJ, Liu L-N. 2020. Decoding the stoichiometric composition and organisation of bacterial metabolosomes. 1. Nature Communications 11:1976.

26. Sutter M, Greber B, Aussignargues C, Kerfeld CA. 2017. Assembly principles and structure of a 6.5-MDa bacterial microcompartment shell. Science 356:1293–1297.

27. Kennedy NW, Ikonomova SP, Slininger Lee M, Raeder HW, Tullman-Ercek D. 2020. Self-assembling shell proteins PduA and PduJ have essential and redundant roles in bacterial microcompartment assembly. Journal of Molecular Biology https://doi.org/10.1016/j.jmb.2020.11.020.

28. Pang A, Frank S, Brown I, Warren MJ, Pickersgill RW. 2014. Structural Insights into Higher Order Assembly and Function of the Bacterial Microcompartment Protein PduA. Journal of Biological Chemistry 289:22377–22384.

29. Fan C, Bobik TA. 2011. The N-terminal region of the medium subunit (PduD) packages adenosylcobalamin-dependent diol dehydratase (PduCDE) into the Pdu microcompartment. Journal of bacteriology 193:5623–5628.

30. Fan C, Cheng S, Liu Y, Escobar CM, Crowley CS, Jefferson RE, Yeates TO, Bobik TA. 2010. Short N-terminal sequences package proteins into bacterial microcompartments. Proceedings of the National Academy of Sciences 107:7509– 7514.

31. Liu Y, Jorda J, Yeates TO, Bobik TA. 2015. The PduL Phosphotransacylase Is Used To Recycle Coenzyme A within the Pdu Microcompartment. Journal of Bacteriology 197:2392–2399.

32. Cheng S, Fan C, Sinha S, Bobik TA. 2012. The PduQ Enzyme Is an Alcohol Dehydrogenase Used to Recycle NAD+ Internally within the Pdu Microcompartment of Salmonella enterica. PLoS ONE 7:e47144.

33. Kerfeld CA, Erbilgin O. 2015. Bacterial microcompartments and the modular construction of microbial metabolism. Trends in Microbiology 23:22–34.

34. Sampson EM, Johnson CLV, Bobik TA. 2005. Biochemical evidence that the pduS gene encodes a bifunctional cobalamin reductase. Microbiology 151:1169–1177.

35. Honda S, Toraya T, Fukui S. 1980. In situ reactivation of glycerol-inactivated coenzyme B12-dependent enzymes, glycerol dehydratase and diol dehydratase. Journal of bacteriology 143:1458–1465.

36. Toraya T, Mori K. 1999. A Reactivating Factor for Coenzyme B12-dependent Diol Dehydratase. Journal of Biological Chemistry 274:3372–3377.

37. Walter D, Ailion M, Roth J. 1997. Genetic characterization of the pdu operon: use of 1, 2-propanediol in Salmonella typhimurium. Journal of bacteriology 179:1013–1022.

38. Chen P, Andersson DI, Roth JR. 1994. The control region of the pdu/cob regulon in Salmonella typhimurium. Journal of bacteriology 176:5474–5482.

39. Bobik TA, Ailion M, Roth JR. 1992. A single regulatory gene integrates control of vitamin B12 synthesis and propanediol degradation. Journal of bacteriology 174:2253–2266.

40. Rondon MR, Escalante-Semerena JC. 1992. The poc locus is required for 1, 2- propanediol-dependent transcription of the cobalamin biosynthetic (cob) and propanediol utilization (pdu) genes of Salmonella typhimurium. Journal of bacteriology 174:2267–2272.

41. Kim EY, Jakobson CM, Tullman-Ercek D. 2014. Engineering Transcriptional Regulation to Control Pdu Microcompartment Formation. PLOS ONE 9:e113814.

42. Nichols TM, Kennedy NW, Tullman-Ercek D. 2020. A genomic integration platform for heterologous cargo encapsulation in 1,2-propanediol utilization bacterial microcompartments. Biochemical Engineering Journal 156:107496.

43. Nichols TM, Kennedy NW, Tullman-Ercek D. 2019. Cargo encapsulation in bacterial microcompartments: Methods and analysis. Methods Enzymol 617:155–186.

44. Sinha S, Cheng S, Fan C, Bobik TA. 2012. The PduM Protein Is a Structural Component of the Microcompartments Involved in Coenzyme B12-Dependent 1,2- Propanediol Degradation by Salmonella enterica. J Bacteriol 194:1912–1918.

45. Parsons JB, Frank S, Bhella D, Liang M, Prentice MB, Mulvihill DP, Warren MJ. 2010. Synthesis of Empty Bacterial Microcompartments, Directed Organelle Protein Incorporation, and Evidence of Filament-Associated Organelle Movement. Molecular Cell 38:305–315.

46. Li Y, Kennedy NW, Li S, Mills CE, Tullman-Ercek D, Olvera de la Cruz M. 2021. Computational and Experimental Approaches to Controlling Bacterial Microcompartment Assembly. ACS Cent Sci 7:658–670.

47. Greber BJ, Sutter M, Kerfeld CA. 2019. The Plasticity of Molecular Interactions Governs Bacterial Microcompartment Shell Assembly. Structure 27:749–763.e4.

48. Chowdhury C, Chun S, Pang A, Sawaya MR, Sinha S, Yeates TO, Bobik TA. 2015. Selective molecular transport through the protein shell of a bacterial microcompartment organelle. Proceedings of the National Academy of Sciences 112:2990–2995.

49. Chowdhury C, Chun S, Sawaya MR, Yeates TO, Bobik TA. 2016. The function of the PduJ microcompartment shell protein is determined by the genomic position of its encoding gene. Mol Microbiol 101:770–783.

50. Mohajerani F, Sayer E, Neil C, Inlow K, Hagan MF. 2021. Mechanisms of Scaffold- Mediated Microcompartment Assembly and Size Control. ACS Nano 15:4197–4212.

51. Engler C, Gruetzner R, Kandzia R, Marillonnet S. 2009. Golden Gate Shuffling: A One-Pot DNA Shuffling Method Based on Type IIs Restriction Enzymes. PLoS ONE 4:e5553.

52. Datta S, Costantino N, Court DL. 2006. A set of recombineering plasmids for gram- negative bacteria. Gene 379:109–115.

53. Schneider CA, Rasband WS, Eliceiri KW. 2012. NIH Image to ImageJ: 25 years of image analysis. Nat Meth 9:671–675.

54. Virtanen P, Gommers R, Oliphant TE, Haberland M, Reddy T, Cournapeau D, Burovski E, Peterson P, Weckesser W, Bright J, van der Walt SJ, Brett M, Wilson J, Millman KJ, Mayorov N, Nelson ARJ, Jones E, Kern R, Larson E, Carey CJ, Polat İ, Feng Y, Moore EW, VanderPlas J, Laxalde D, Perktold J, Cimrman R, Henriksen I, Quintero EA, Harris CR, Archibald AM, Ribeiro AH, Pedregosa F, van Mulbregt P. 2020. SciPy 1.0: fundamental algorithms for scientific computing in Python. Nat Methods 17:261–272.

55. Crowley CS, Cascio D, Sawaya MR, Kopstein JS, Bobik TA, Yeates TO. 2010. Structural insight into the mechanisms of transport across the Salmonella enterica Pdu microcompartment shell. Journal of Biological Chemistry 285:37838–37846.

56. Pang A, Liang M, Prentice MB, Pickersgill RW. 2012. Substrate channels revealed in the trimeric Lactobacillus reuteri bacterial microcompartment shell protein PduB. Acta Cryst D, Acta Cryst Sect D, Acta Crystallogr D, Acta Crystallogr Sect D, Acta Crystallogr D Biol Crystallogr, Acta Crystallogr Sect D Biol Crystallogr 68:1642– 1652.

57. Pettersen EF, Goddard TD, Huang CC, Couch GS, Greenblatt DM, Meng EC, Ferrin TE. 2004. UCSF Chimera--a visualization system for exploratory research and analysis. J Comput Chem 25:1605–1612.

