## Supplementary Document 1 for "Linking the *Salmonella enterica* 1,2-propanediol utilization bacterial microcompartment shell to the enzymatic core via the shell protein PduB"

#### 1 Introduction

The kinetic model for the 1,2-propanediol utilization (Pdu) pathway here was developed based on the original work of Jakobson *et al.*[1] Specifically, we assume in this model that the cytosol is well-mixed, such that we are working with a compartmental model. We also incorporate the effects of cell growth over time, and explicitly model the metabolite concentrations in the external media. This required several adjustments to the original model, outlined below. We focus our analysis here on metabolite profiles over time in a batch reactor setting instead of at steady-state, to study the conditions under which we ran our growth curves.

#### 2 Model Assumptions

We assume the following in our model:

1. At time $t$, there are $N(t)$ identical, non-interacting cells in a well mixed solution.
2. The substrates 1,2-propanediol, propionaldehyde, propionyl-CoA, propionate, and 1-propanol passively diffuse across the cell membrane at rates specified by permeability parameters.
3. The substrates 1,2-propanediol, propionaldehyde, propionyl-CoA, propionate, and 1-propanol passively diffuse across the microcompartment (MCP) shell/polar body surface at rates specified by permeability parameters.
4. There are $n_{\text{MCP}}$ non-interacting MCPs or polar bodies in the cytosol of each cell.
5. Reactions catalyzed by PduCDE, PduP, and PduQ, forward or reverse, can only occur in the MCP or polar body
6. Reactions catalyzed by PduL/W can only occur in the cytosol.
7. The external media, cytosol of the cell, and internal MCP volume are well-mixed such that the concentration in each compartment is assumed uniform at any given point in time.
8. The volume of the external media up to leading order is the volume of the entire culture.
9. The volume of the cytosol up to leading order is the volume of the cell.
10. All enzymes behave according to Michaelis-Menten kinetics.

### 3 Chemical Reactions

The following reactions are considered in our model, with the assumptions described above:

$$\text{1,2-propanediol} \xrightarrow{\text{PduCDE}} \text{Propionaldehyde}$$

$$\text{propionaldehyde} \xleftrightarrow{\text{PduQ}} \text{1-propanol}$$

$$\text{propionaldehyde} \xleftrightarrow{\text{PduP}} \text{propionyl-CoA}$$

$$\text{propionyl-CoA+} \xrightarrow{\text{PduLW}} \text{propionate}$$

### 4 Equations Used In Mathematical Model

The differential equations described below were integrated forward in time from a starting condition of 55 mM 1,2-propanediol in the external media, the starting condition for our growth curve.

#### 4.1 Variable Definition

Concentrations of all substrates are defined by the following variables:

- $P_i$: 1,2-propanediol concentration in volume $i$ (MCP/polar body, cytosol, or external media)
- $A_i$: propionaldehyde concentration in volume $i$ (MCP/polar body, cytosol, or external media)
- $Pol_i$: 1-propanol concentration in volume $i$ (MCP/polar body, cytosol, or external media)
- $PCoA_i$: propionyl-CoA concentration in volume $i$ (MCP/polar body, cytosol, or external media)
- $Pate_i$: propionate concentration in volume $i$ (MCP/polar body, cytosol, or external media)

Other constants and variables used in the model are defined follows:

- $V_{\text{compartment}}$: volume of the compartment (MCP or polar body)
- $SA_{\text{compartment}}$: surface area of the compartment (MCP or polar body)
- $Vol_{\text{cell}}$: volume of the cell
- $SA_{\text{cell}}$: surface area of the cell
- $Perm_i^j$: permeability of substrate $i$ at interface $j$ (cell surface or MCP/polar body surface)
- $R_i(X_j)$: reaction rate of enzyme $i$ as a function of concentrations $X$ in a given volume $j$ (MCP/polar body, cytosol, or external media)
- $K_{\text{M}}^{i,j}$: Michaelis constant of enzyme $i$ for substrate $j$
- $V_{\text{max}}^{i,j}$: Maximum reaction velocity of enzyme $i$ for substrate $j$
- $N(t)$: Number of cells at time $t$, calculated from the experimental growth profile.
- PB: Refers to polar body
- MCP: Refers to MCP
- $n_{\text{MCP}}$: Number of MCPs/polar bodies per cell

#### 4.2 Differential Equations: MCP or Polar Body

The differential equations for the MCP or polar body volume were as follows:

$$
\begin{aligned}
\frac{dP_{\text{MCP/PB}}}{dt} =& -R_{\text{PduCDE}}\left(X_{\text{MCP/PB}}\right) + \frac{Perm^{\text{P}}_{\text{MCP/PB}} SA_{\text{MCP/PB}}}{Vol_{\text{MCP/PB}}}\left(P_{\text{cytosol}} - P_{\text{MCP/PB}}\right) \\
\frac{dA_{\text{MCP/PB}}}{dt} =& R_{\text{PduCDE}}\left(X_{\text{MCP/PB}}\right) - R_{\text{PduP},f}\left(X_{\text{MCP/PB}}\right) + R_{\text{PduP},r}\left(X_{\text{MCP/PB}}\right) \\
& - R_{\text{PduQ},f}\left(X_{\text{MCP/PB}}\right) + R_{\text{PduQ},r}\left(X_{\text{MCP/PB}}\right) \\
& + \frac{Perm^{\text{A}}_{\text{MCP/PB}} SA_{\text{MCP/PB}}}{Vol_{\text{MCP/PB}}}\left(A_{\text{cytosol}} - A_{\text{MCP/PB}}\right) \\
\frac{dPol_{\text{MCP/PB}}}{dt} =& R_{\text{PduQ},f}\left(X_{\text{MCP/PB}}\right) - R_{\text{PduQ},r}\left(X_{\text{MCP/PB}}\right) \\
& + \frac{Perm^{\text{Pol}}_{\text{MCP/PB}} SA_{\text{MCP/PB}}}{Vol_{\text{MCP/PB}}}\left(Pol_{\text{cytosol}} - Pol_{\text{MCP/PB}}\right) \\
\frac{dPCoA_{\text{MCP/PB}}}{dt} =& R_{\text{PduP},f}\left(X_{\text{MCP/PB}}\right) - R_{\text{PduP},r}\left(X_{\text{MCP/PB}}\right) \\
& + \frac{Perm^{\text{PCoA}}_{\text{MCP/PB}} SA_{\text{MCP/PB}}}{Vol_{\text{MCP/PB}}}\left(PCoA_{\text{cytosol}} - PCoA_{\text{MCP/PB}}\right) \\
\frac{dPate_{\text{MCP/PB}}}{dt} =& \frac{Perm^{\text{Pate}}_{\text{MCP/PB}} SA_{\text{MCP/PB}}}{Vol_{\text{MCP/PB}}}\left(Pate_{\text{cytosol}} - Pate_{\text{MCP/PB}}\right)
\end{aligned}
$$

#### 4.3 Differential Equations: Cytosol

The differential equations for the cytosol of the cell were as follows:

$$\begin{aligned}
\frac{dP_{\text{cytosol}}}{dt} =& \frac{n_{\text{MCP}} Perm^{\text{P}}_{\text{MCP/PB}} SA_{\text{MCP/PB}}}{Vol_{\text{MCP/PB}}} \left(P_{\text{MCP/PB}} - P_{\text{cytosol}}\right) \\
&+ \frac{Perm^{\text{P}}_{\text{cell}} SA_{\text{cell}}}{Vol_{\text{cell}}} \left(P_{\text{external}} - P_{\text{cytosol}}\right) \\
\frac{dA_{\text{cytosol}}}{dt} =& \frac{n_{\text{MCP}} Perm^{\text{A}}_{\text{MCP/PB}} SA_{\text{MCP/PB}}}{Vol_{\text{MCP/PB}}} \left(A_{\text{MCP/PB}} - A_{\text{cytosol}}\right) \\
&+ \frac{Perm^{\text{A}}_{\text{cell}} SA_{\text{cell}}}{Vol_{\text{cell}}} \left(A_{\text{external}} - A_{\text{cytosol}}\right) \\
\frac{dPol_{\text{cytosol}}}{dt} =& \frac{n_{\text{MCP}} Perm^{\text{Pol}}_{\text{MCP/PB}} SA_{\text{MCP/PB}}}{Vol_{\text{MCP/PB}}} \left(Pol_{\text{MCP/PB}} - Pol_{\text{cytosol}}\right) \\
&+ \frac{Perm^{\text{Pol}}_{\text{cell}} SA_{\text{cell}}}{Vol_{\text{cell}}} \left(Pol_{\text{external}} - Pol_{\text{cytosol}}\right) \\
\frac{dPCoA_{\text{cytosol}}}{dt} =& - R_{\text{PduLW}} \left(X_{\text{cytosol}}\right) \\
&+ \frac{n_{\text{MCPB}} Perm^{\text{PCoA}}_{\text{MCP/PB}} SA_{\text{MCP/PB}}}{Vol_{\text{MCP/PB}}} \left(PCoA_{\text{MCP/PB}} - PCoA_{\text{cytosol}}\right) \\
&+ \frac{Perm^{\text{PCoA}}_{\text{cell}} SA_{\text{cell}}}{Vol_{\text{cell}}} \left(PCoA_{\text{external}} - PCoA_{\text{cytosol}}\right) \\
\frac{dPate_{\text{cytosol}}}{dt} =& R_{\text{PduLW}} \left(X_{\text{cytosol}}\right) \\
&+ \frac{n_{\text{MCP}} Perm^{\text{Pate}}_{\text{MCP/PB}} SA_{\text{MCP/PB}}}{Vol_{\text{MCP/PB}}} \left(Pate_{\text{MCP/PB}} - Pate_{\text{cytosol}}\right) \\
&+ \frac{Perm^{\text{Pate}}_{\text{cell}} SA_{\text{cell}}}{Vol_{\text{cell}}} \left(Pate_{\text{external}} - Pate_{\text{cytosol}}\right)
\end{aligned}$$

#### 4.4 Differential Equations: External Media

The differential equations for the external media of the cell were as follows:

$$\begin{aligned}
\frac{dP_{\text{external}}}{dt} =& N(t) \frac{Perm^{\text{P}}_{\text{cell}} SA_{\text{cell}}}{Vol_{\text{cell}}} \left(P_{\text{external}} - P_{\text{cytosol}}\right) \\
\frac{dA_{\text{external}}}{dt} =& N(t) \frac{Perm^{\text{A}}_{\text{cytosol}} SA_{\text{cell}}}{Vol_{\text{cell}}} \left(A_{\text{external}} - A_{\text{cytosol}}\right) \\
\frac{dPol_{\text{external}}}{dt} =& N(t) \frac{Perm^{\text{Pol}}_{\text{cytosol}} SA_{\text{cell}}}{Vol_{\text{cell}}} \left(Pol_{\text{external}} - Pol_{\text{cytosol}}\right) \\
\frac{dPCoA_{\text{external}}}{dt} =& N(t) \frac{Perm^{\text{PCoA}}_{\text{cytosol}} SA_{\text{cell}}}{Vol_{\text{cell}}} \left(PCoA_{\text{external}} - PCoA_{\text{cytosol}}\right) \\
\frac{dPate_{\text{external}}}{dt} =& N(t) \frac{Perm^{\text{Pate}}_{\text{cytosol}} SA_{\text{cell}}}{Vol_{\text{cell}}} \left(Pate_{\text{external}} - Pate_{\text{cytosol}}\right)
\end{aligned}$$

#### 4.5 Definition of Reaction Rate

Reaction rates were assumed to follow Michaelis-Menten kinetics

$$R_{\text{PduCDE}} = V_{\text{max}}^{\text{PduCDE}} \frac{P_{\text{MCP/PB}}}{K_{\text{M}}^{\text{PduCDE}} + P_{\text{MCP/PB}}}$$
$$R_{\text{PduP},f} = V_{\text{max}}^{\text{PduP},f} \frac{A_{\text{MCP/PB}}}{K_{\text{M}}^{\text{PduP,f}} + A_{\text{MCP/PB}}}$$
$$R_{\text{PduP},r} = V_{\text{max}}^{\text{PduP},r} \frac{PCoA_{\text{MCP/PB}}}{K_{\text{M}}^{\text{PduP,r}} + PCoA_{\text{MCP/PB}}}$$
$$R_{\text{PduQ},f} = V_{\text{max}}^{\text{PduQ},f} \frac{A_{\text{MCP/PB}}}{K_{\text{M}}^{\text{PduQ,f}} + A_{\text{MCP/PB}}}$$
$$R_{\text{PduQ},r} = V_{\text{max}}^{\text{PduQ},r} \frac{Pol_{\text{MCP/PB}}}{K_{\text{M}}^{\text{PduQ,r}} + Pol_{\text{MCP/PB}}}$$
$$R_{\text{PduLW}} = V_{\text{max}}^{\text{PduLW}} \frac{PCoA_{\text{cytosol}}}{K_{\text{M}}^{\text{PduLW}} + PCoA_{\text{cytosol}}}$$

### References

[1] Christopher M. Jakobson, Danielle Tullman-Ercek, Marilyn F. Slininger, and Niall M. Mangan. A systems-level model reveals that 1,2-propanediol utilization microcompartments enhance pathway flux through intermediate sequestration. *PLOS Computational Biology*, 13(5):1–24, 05 2017.
