## Supplemental Tables and Figures for "Linking the *Salmonella enterica* 1,2-propanediol utilization bacterial microcompartment shell to the enzymatic core via the shell protein PduB"

**Table S1.** Strains used in this study.

| **Strain** | **Organism** | **Genotype** |
| --- | --- | --- |
| DTE003 | *S. enterica* serovar Typhimurium LT2 | Wild type |
| CMJS256 | *S. enterica* serovar Typhimurium LT2 | Δ*pocR* |
| TUC01 [1], [2] | *E. coli* W3110 | *gal490 pglΔ8 λcI857 Δ(cro‐bioA) int<>cat/sacB* |
| NWKs083 | *S. enterica* serovar Typhimurium LT2 | Δ*pduA* Δ*pduJ* |
| CEMS179 | *S. enterica* serovar Typhimurium LT2 | Δ*pduB* |
| NWKs365 | *S. enterica* serovar Typhimurium LT2 | Δ*pduB::pduB-K102A-K207A* |
| NWKs337 | *S. enterica* serovar Typhimurium LT2 | Δ*pduA::pduA-mCherry* |
| NWKs338 | *S. enterica* serovar Typhimurium LT2 | Δ*pduA::pduA-mCherry* Δ*pduB* |
| NWKs344 | *S. enterica* serovar Typhimurium LT2 | Δ*pduA::pduA-mCherry* Δ*pocR* |
| NWKs364 | *S. enterica* serovar Typhimurium LT2 | Δ*pduA::pduA-mCherry* Δ*pduB::pduB-K102A-K207A* |
| TMDS025 | *S. enterica* serovar Typhimurium LT2 | Δ*pduD::ssD-GFPmut2* |
| CEMS299 | *S. enterica* serovar Typhimurium LT2 | Δ*pocR* Δ*pduD::ssD-GFPmut2* |
| CEMS298 | *S. enterica* serovar Typhimurium LT2 | Δ*pduA* Δ*pduD::ssD-GFPmut2* Δ*pduJ* |
| CEMS218 | *S. enterica* serovar Typhimurium LT2 | Δ*pduB* Δ*pduD::ssD-GFPmut2* |
| CEMS296 | *S. enterica* serovar Typhimurium LT2 | Δ*pduB::pduB-K102A-K207A* Δ*pduD::ssD-GFPmut2* |

**Table S2.** Plasmids used in this study.

| **Name** | **Plasmid** | **Origin** | **Resistance** |
| --- | --- | --- | --- |
| CMJ069 | pBAD33t-ssD-GFPmut2 | p15A | Chloramphenicol |
| EYK208 | pBAD33t-ssP-GFPmut2 | p15A | Chloramphenicol |
| EYK193 | pBAD33t-PduG-GFPmut2 | p15A | Chloramphenicol |
| NWKp041 | pBAD33t-PduO-GFPmut2 | p15A | Chloramphenicol |
| NWKp042 | pBAD33t-PduW-GFPmut2 | p15A | Chloramphenicol |
| NWKp043 | pBAD33t-ssL-GFPmut2 | p15A | Chloramphenicol |
| pSIM6 [2] | λ Red system repressed by cI857 | pSC101 *repA^ts^* | Ampicillin |

**Table S3.** Primers used in this study.

| **Name** | **Purpose** | **Description** | **Sequence*** |
| --- | --- | --- | --- |
| NWKo493 | Recombineering | Amplify *cat/sacB* with homology upstream of *pduB* For | gccctcacaccgatgtagaaaaaatcttaccgaagggaattagccaatgaTGTGACGGAAGATCACTTCG |
| NWKo494 | Recombineering | Amplify *cat/sacB* with homology downstream of *pduB* Rev | ttcgccagtgcttcaaatcttttcgatctcatgaatcagcctcgtgggtaATCAAAGGGAAAACTGTCCATAT |
| NWKo517 | Recombineering | Amplify *pduB-K102A-K207A* with homology upstream of *pduB* For | ccacgccctcacaccgatgtagaaaaaatcttaccgaagggaattagccaATGAGCAGCAATGAGCTGG |
| NWKo518 | Recombineering | Amplify *pduB-K102A-K207A* with homology downstream of *pduB* Rev | gccagtgcttcaaatcttttcgatctcatgaatcagcctcgtgggtatcaGATGTAGGACGGACGATCG |
| NWKo499 | Recombineering | Knockout *pduB* | cgcgtccgcacagcgatgttgaggccattttaccgaaatcagcctaatcgggcaagcgaggtgaagcgtaatggataaagagcttctgcaatcaacggtc |
| NWKo466 | Sequencing | Amplify from upstream of *pduA* | CTGCGAACCTGTCTCC |
| NWKo467 | Sequencing | Amplify from downstream of *pduA* | CGTCTCTCGTATAGGTTGG |
| NWKo501 | Golden Gate cloning | Amplify *pduO* with GG overhang For | AttGGTCTCACATGGCGATTTATACCCGAAC |
| NWKo504 | Golden Gate cloning | Amplify *pduO* with GG overhang and GS linker Rev | ATTGGTCTCAGCTGCCttgatgagttcccacgttaatag |
| NWKo502 | Golden Gate cloning | Amplify *pduW* with GG overhang For | AttGGTCTCACATGTCTTACAAAATAATGGCCATTA |
| NWKo505 | Golden Gate cloning | Amplify *pduW* with GG overhang and GS linker Rev | ATTGGTCTCAGCTGCCggctggtacacaaagcc |
| EYKP263 | SacI cloning | Amplify *pduG* with SacI recognition For | TATTGAGCTCTTAAAGAGGAGAAAGGTCatgcgatatatagctggcattgacatc |
| EYKP264 | XbaI cloning | Amplify *pduG* with XbaI recognition Rev | TATTTCTAGActgtccatgcgcaaactccttatg |
| NWKo506 | Golden Gate cloning | Amplify *GFPmut2* with GG overhang and GS linker For | AttGGTCTCACAGCAgtaaaggagaagaacttttcactgg |
| NWKo507 | Golden Gate cloning | Amplify *GFPmut2* with GG overhang Rev | ATTGGTCTCATTTAtttgtatagttcatccatgccatgtg |
| TMDP058 | Golden Gate cloning | Amplify *ssL-GFPmut2* with GG overhang For | ATTGGTCTCACATGgataaagagcttctgcaatcaac |
| NWKo514 | QuikChange | *PduB-K102A* For | ATGGCGGCGGACGAAGCGGTGGCGGCCACCAACACC |
| NWKo515 | QuikChange | *PduB-K102A* Rev | CTGCTTCGCCACCGCCGGTGGTTGTGGCTTCACCAG |
| NWKo516 | QuikChange | *PduB-K207A* For | ATACCGCGCTGGCGTCAGCCAACGTTGAAGTCG |
| NWKo517 | QuikChange | *PduB-K207A* Rev | ACTACCGGCTATGGCGCGACCGCAGTCGGTTGC |
| MPV031 | Sequencing | Sequence confirmation of *pduB* locus For | gccctcacaccgatgtagaaaaaatc |
| MPV030 | Sequencing | Sequence confirmation of *pduB* locus Rev | tccatcgccatcacttcttcg |
| TMDP021 | Sequencing, recombineering | Amplify from upstream of *pduD* locus For | ggcaacaggttatcgcctgc |
| TMDP022 | Sequencing, recombineering | Amplify from downstream of *pduD* locus Rev | ccctgcaggctgttcatgc |
| TMDP072 | Sequencing | Amplify from upstream of *pduD* locus For | catccagaaagccaagctaacc |
| TMDP073 | Sequencing | Amplify from downstream of *pduD* locus Rev | cgtccagcgttttattggtgg |
| TMDP016 | Recombineering | Amplify ssD-GFPmut2 with homology upstream of *pduD* For | TTGATCCCAACGAGATTGATTAAGGGGTGAGAAatggaaattaatgaaaaattgctgcgc |
| TMDP019 | Recombineering | Amplify ssD-GFPmut2 with homology downstream of *pduD* Rev | ATTGCGTCGGTATTCATGGAGTTATCCTTTAttatttgtatagttcatccatgccatgtg |

**Table S4.** Parameter values used in kinetic model of pathway performance in MCPs and polar bodies.

| **Parameter Name** | **MCP** | **Polar Body** | **Unit** |
| --- | --- | --- | --- |
| CDE_con | 0.462467024 | 0.462467024 | mM |
| CDE_tot [3] | 6000 | 6000 | enzymes/cell |
| cell_length | 2.47E-06 | 2.47E-06 | m |
| cell_radius | 3.75E-07 | 3.75E-07 | m |
| cell_surface_area | 5.82E-12 | 5.82E-12 | m^2^ |
| cell_volume | 9.81E-19 | 9.81E-19 | m^3^ |
| external_volume | 5.00E-05 | 5.00E-05 | m^3^ |
| kcatCDE [4] | 300 | 300 | 1/s |
| kcatL | 100 | 100 | 1/s |
| kcatPf [5] | 55 | 55 | 1/s |
| kcatPr [5] | 6 | 6 | 1/s |
| kcatQf [6] | 55 | 55 | 1/s |
| kcatQr [6] | 6 | 6 | 1/s |
| L_con [7] | 0.1 | 0.1 | mM |
| mcp_surface_area | 6.16E-14 | 3.75E-13 | m^2^/compartment |
| mcp_volume | 1.44E-21 | 2.16E-20 | m^3^/compartment |
| Navogadro | 6.02E+23 | 6.02E+23 | molecules/mole |
| nmcp | 15 | 1 | MCPs/polar bodies per cell |
| P_con | 0.693700536 | 0.693700536 | mM |
| P_tot [3] | 9000 | 9000 | enzymes/cell |
| PermMCPNonPolar | 3.98E-08 | 1.00E+01 | m/s |
| PermMCPPolar | 3.98E-08 | 1.00E+01 | m/s |
| Q_con | 0.520275402 | 0.520275402 | mM |
| Q_tot [3] | 6750 | 6750 | enzymes/cell |
| radius_mcp [8] | 7.00E-08 | 1.73E-07 | m |
| VmaxCDEf | 138.7401071 | 138.7401071 | mM/s |
| VmaxLf | 10 | 10 | mM/s |
| VmaxPf | 38.15352946 | 38.15352946 | mM/s |
| VmaxPr | 4.162203213 | 4.162203213 | mM/s |
| VmaxQf | 28.61514709 | 28.61514709 | mM/s |
| VmaxQr | 3.12165241 | 3.12165241 | mM/s |
| KmCDEPropanediol [4] | 0.5 | 0.5 | mM |
| KmPfPropionaldehyde [5] | 15 | 15 | mM |
| KmPrPropionyl [5] | 95 | 95 | mM |
| KmQfPropionaldehyde [6] | 15 | 15 | mM |
| KmQrPropanol [6] | 95 | 95 | mM |
| KmLPropionyl | 20 | 20 | mM |
| PermCellPropanediol [9] [10] [11] | 1.00E-04 | 1.00E-04 | m/s |
| PermCellPropionaldehyde [9] [10] [11] | 1.00E-02 | 1.00E-02 | m/s |
| PermCellPropanol [9] [10] [11] | 1.00E-04 | 1.00E-04 | m/s |
| PermCellPropionyl [9] [10] [11] | 1.00E-05 | 1.00E-05 | m/s |
| PermCellPropionate [9] [10] [11] | 1.00E-07 | 1.00E-07 | m/s |

**Figure S1.** Tornado plot showing changes in maximum propionaldehyde level observed in MCP model with a 10% increase and decrease in the noted parameter.

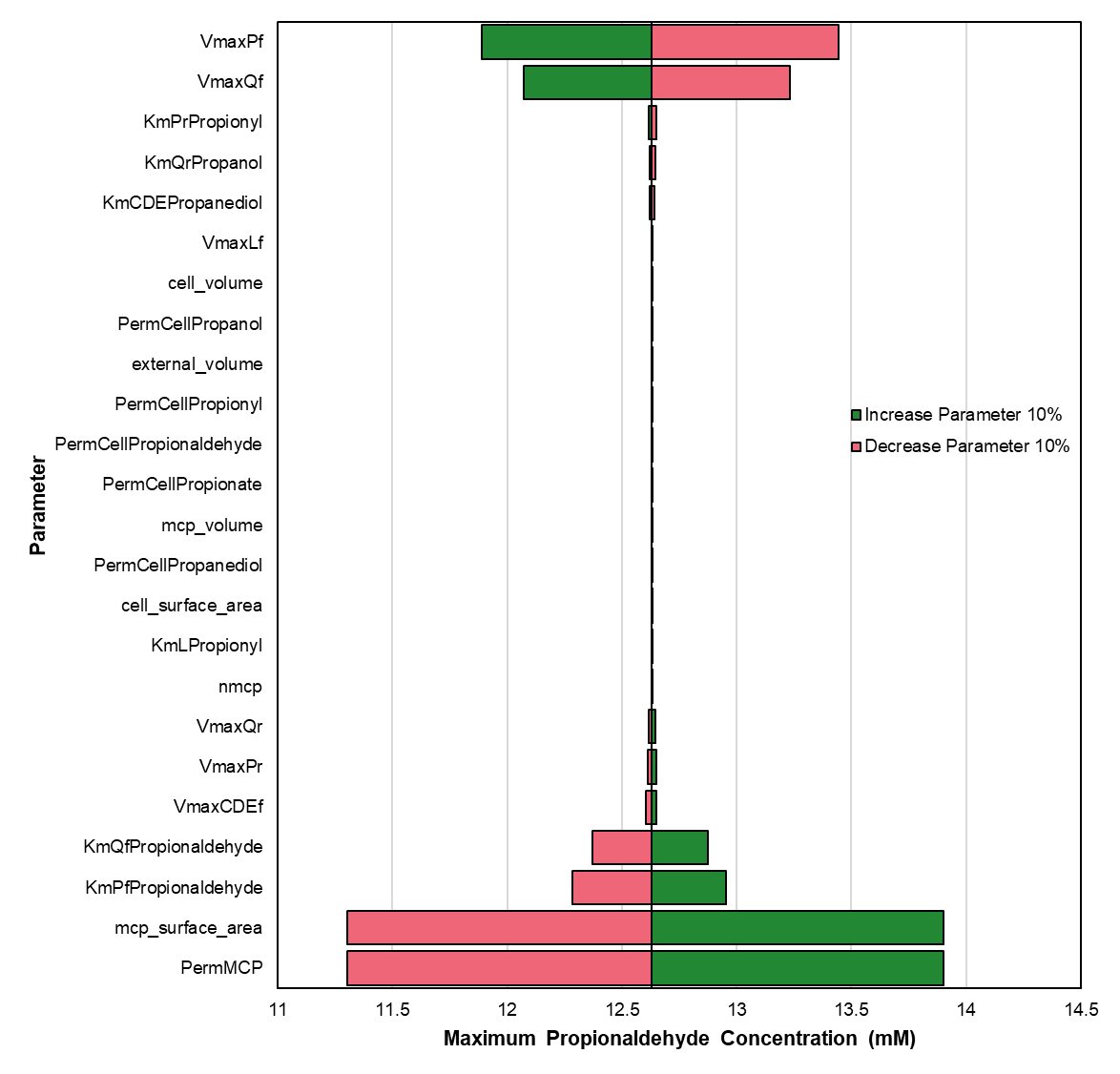

**Figure S2.** Tornado plot showing changes in maximum propionaldehyde level observed in polar body model with a 10% increase and decrease in the noted parameter.

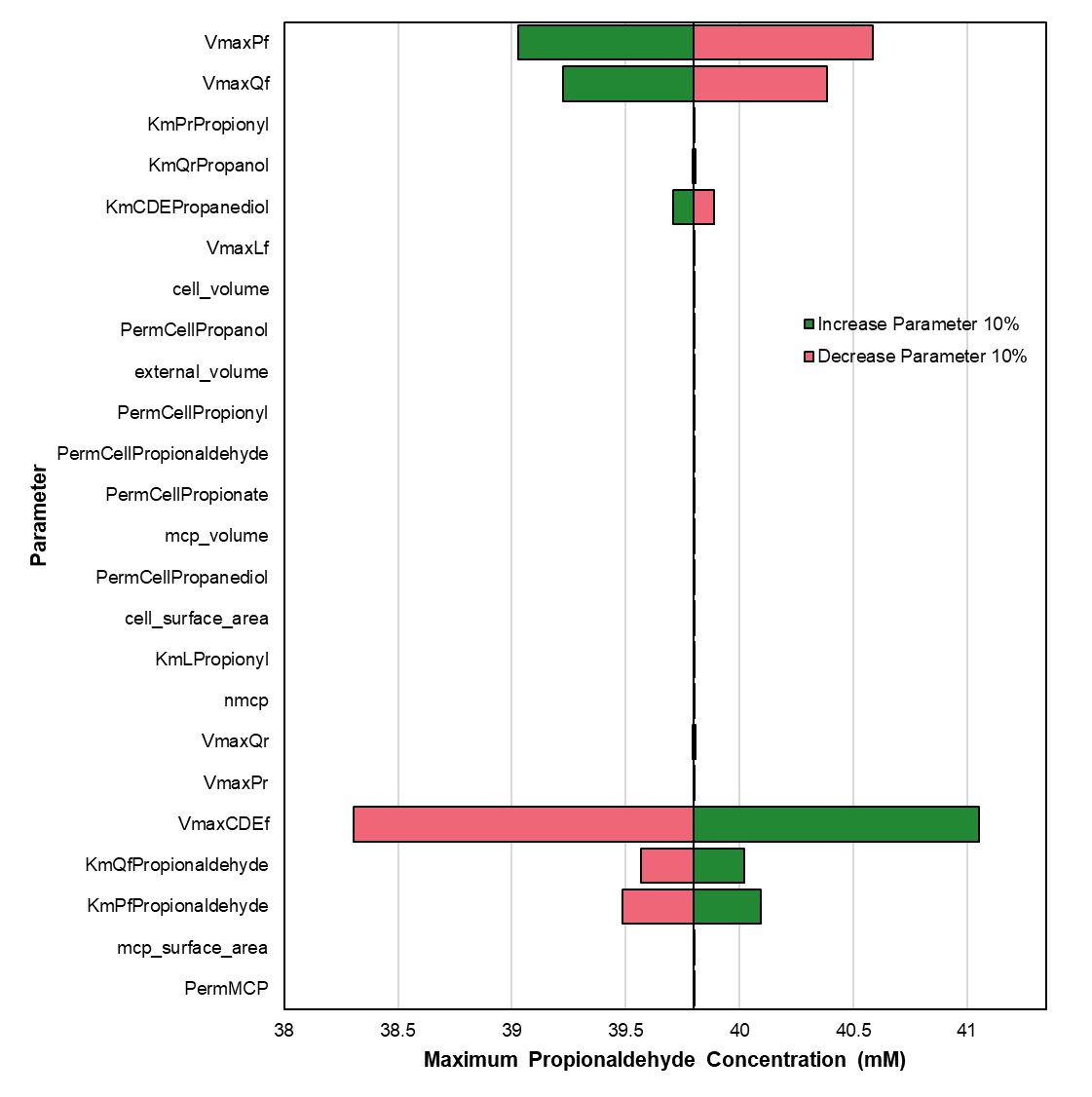

**Figure S3.** Full profiles of all metabolites for the kinetic model results shown in Figure 7. Profiles are shown for the (a) MCP model with 0.46 mM PduCDE and the polar body model with (b) 0.46 mM PduCDE, (c) 0.23 mM PduCDE, and (d) 0.12 mM PduCDE.

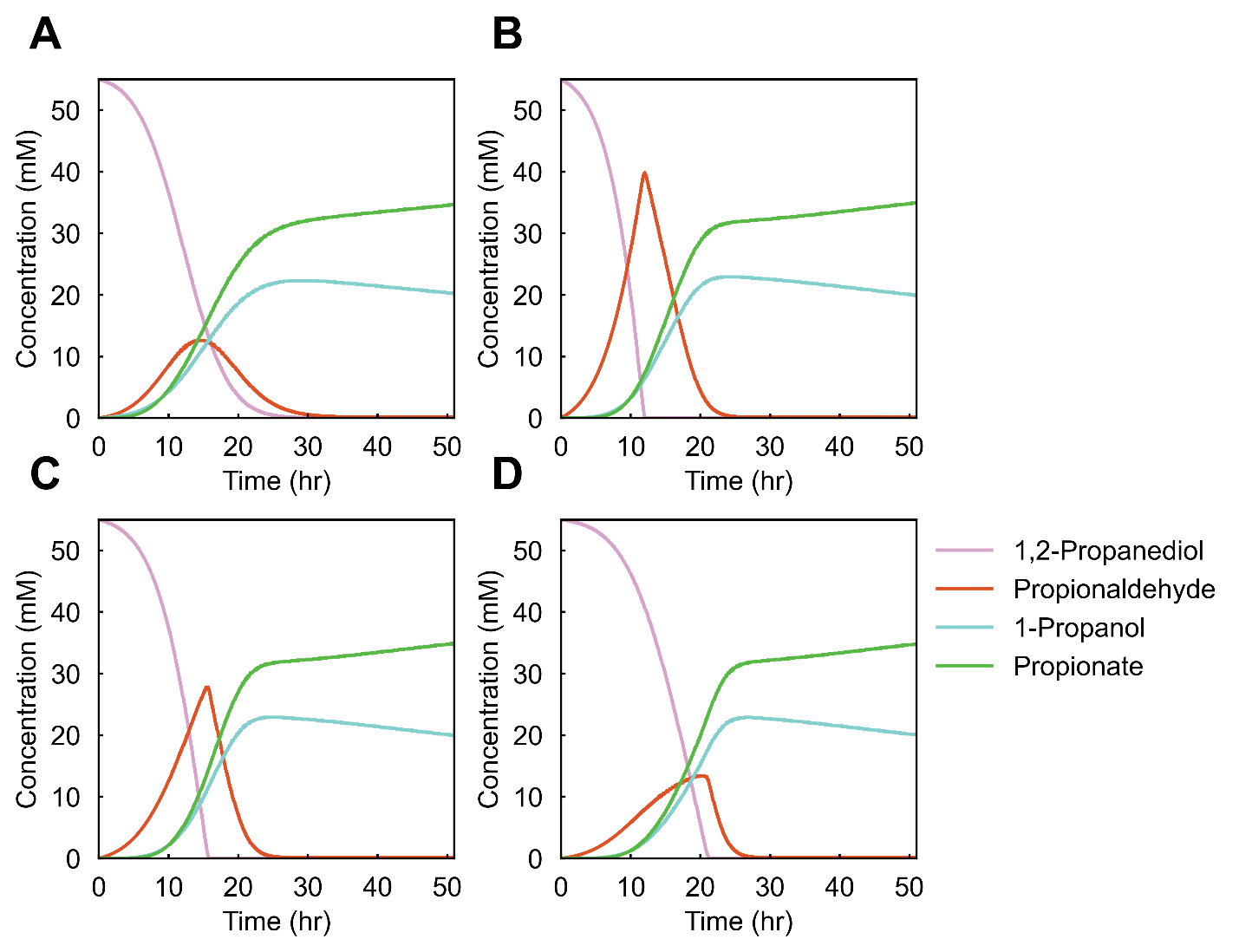

**Bibliography**

[1]	L.C. Thomason, J.A. Sawitzke, X. Li, N. Costantino, D.L. Court, Recombineering: genetic engineering in bacteria using homologous recombination, Curr. Protoc. Mol. Biol. 106 (2014) 1.16.1-1.16.39. https://doi.org/10.1002/0471142727.mb0116s106.

[2]	S. Datta, N. Costantino, D.L. Court, A set of recombineering plasmids for gram-negative bacteria, Gene. 379 (2006) 109–115. https://doi.org/10.1016/j.gene.2006.04.018.

[3]	M. Yang, D.M. Simpson, N. Wenner, P. Brownridge, V. M. Harman, J.C.D. Hinton, R.J. Beynon, L. Liu, Decoding the stoichiometric composition and organisation of bacterial metabolosomes, Nat. Comm. 11 (2020) 1976. https://doi.org/10.1038/s41467-020-15888-4.

[4]	W.W. Bachovichin, R.G. Eagar, Jr., K.W. Moore, J.H. Richards, Mechanism of action of adenosylcobalamin: glycerol and other substrate analogs as substrates and inactivators for propanediol dehydratase - kinetics, stereospecificity, and mechanism, Biochemistry. 16 (1977) 1082-1092. https://doi.org/10.1021/bi00625a009.

[5]	N.A. Leal, G.D. Havemann, T.A. Bobik, PduP is a coenzyme-a-acylating propionaldehyde dehydrogenase associated with the polyhedral bodies involved in B12-dependent 1,2-propanediol degradation by Salmonella enterica serovar Typhimurium LT2, Arch Microbiol. 180 (2003) 353-361. https://doi.org/10.1007/s00203-003-0601-0.

[6]	S. Cheng, C. Fan, S. Sinha, T.A. Bobik, The PduQ Enzyme Is an Alcohol Dehydrogenase Used to Recycle NAD^+^ Internally within the Pdu Microcompartment of *Salmonella enterica*, PLoS ONE. 7 (2012) e47144. https://doi.org/10.1371/journal.pone.0047144.

[7]	K.R. Able, M.H. Butler, B.E.Wright, Cellular Concentrations of Enzymes and Their Substrates, J. theor. Biol. 143 (1990) 163-195.

[8]	N.W. Kennedy, J.M. Hershewe, T.M. Nichols, E.W. Roth, C.D. Wilke, C.E. Mills, M.C. Jewett, D. Tullman-Ercek, Apparent size and morphology of bacterial microcompartments varies with technique, PLos ONE. 15 (2019) e0226395. https://doi.org/10.1371/journal.pone.0226395.

[9]	E. Orbach, A. Finkelstein, The nonelectrolyte permeability of planar lipid bilayer membranes, J. Gen. Physiol. 75 (1980) 427-436.

[10]	M.H. Abraham, G.S. Whiting, R. Fuchs, E.J. Chambers, Thermodynamics of solute transfer from water to hexadecane, J. Chem. Soc., Perkin Trans. 2. (1990) 291-300. https://doi.org/10.1039/P29900000291.

[11]	M.M. Schantz, D.E. Martire, Determination of hydrocarbon-water partition coefficients from chromatographic data and based on solution thermodynamics and theory, J. Chromatogr. A. 391 (1987) 35-51.
